# Improved pathogen and stress tolerance in tomato mutants of SET domain histone 3 lysine methyltransferases

**DOI:** 10.1101/2022.02.19.481141

**Authors:** Carol Bvindi, Sanghun Lee, Liang Tang, Michael V. Mickelbart, Ying Li, Tesfaye Mengiste

## Abstract

Histone lysine methylations (HLMs) are implicated in control of gene expression in different eukaryotes. However, the role of HLMs in regulating desirable crop traits and the enzymes involved in these modifications are poorly understood. We studied the functions of tomato histone H3 lysine methyltransferases Set Domain Group 33 (SDG33) and SDG34 in biotic and abiotic stress responses. *SDG33* and *SDG34* mutants were altered in H3K36 and H3K4 methylations, and expression of genes involved in diverse processes and responses to biotic and abiotic stimuli. The double but not the single mutants show resistance to the fungal pathogen *Botrytis cinerea.* Interestingly, single mutants were tolerant to drought and the double mutant showed superior tolerance consistent with independent and additive functions. Mutants maintained higher water status during drought and improved recovery and survival after lapse of drought. Notably, diminution of H3K4 and H3K36 trimethylation and expression of negative regulators in challenged plants contributes to stress tolerance of the mutants. Mutations in SDG33 and SDG34 remove predisposition to biotic and abiotic stress by disrupting permissive transcriptional context promoting expression of negative regulatory factors. These allows improvement of stress and pathogen tolerance through modification of histone epigenetic marks.

## Introduction

A hierarchy of regulatory factors from perception of extracellular signals to activation and production of protective molecules contribute to plant responses to environmental stress. The perception of environmental cues and response signaling networks allow plants to rapidly reprogram and activate coping mechanisms. Responses to environmental assaults such as drought are mediated by complex molecular mechanisms and genetic regulators that mediate changes in cellular, developmental, and physiological processes (Zhu, 2016). Similarly, sophisticated microbial sensing and signaling mechanisms mediate activation of plant immune responses (Jones & Dangl, 2006). Transcriptional reprogramming of a battery of genes and their spatio-temporal regulation is a major component of plant adaptation to biotic and abiotic stresses (Fujita *et al*., 2006). The details of how transcription reprogramming occurs and the co-regulation of diverse genes at the genome scale in the context of chromatin are not well understood.

In eukaryotes, DNA is wrapped in a nucleoprotein complex known as chromatin that prevents access of the transcriptional machinery to gene regulatory sequences. Histone epigenetic marks modulate such accessibility and provide the chromatin context for transcription (Kouzarides, 2007). Histone Lysine methylation (HLM) is a major histone epigenetic mark catalyzed by histone lysine methyltransferases (HLMTs). Histone lysine residues can be mono-, di, or tri-methylated altering chromatin structure, influencing the accessibility of the transcriptional machinery to the corresponding DNA, and modulating transcription of targeted genes (Kouzarides, 2007). Plant HLMTs carry a conserved <u>S</u>U- (VAR) 3-9, <u>E</u>nhance of zeste E (Z) and <u>T</u>rithorax (SET) domain responsible for their enzymatic activity. Studies in Arabidopsis have implicated HLMTs in regulating plant growth (Dong *et al*., 2008; Xu *et al*., 2008; Cazzonelli *et al*., 2009), flowering time (Zhao *et al*., 2005; Xu *et al*., 2008), and responses to biotic and abiotic stress (Chinnusamy & Zhu, 2009; Pandey *et al*., 2016). Changes in HLM correlating with transcriptional status of target genes in response to pathogens (Berr *et al*., 2010; Lee *et al*., 2016), cold (Hu *et al*., 2012), heat (Pecinka *et al*., 2010; Folsom *et al*., 2014) and salt stress (Song *et al*., 2012) have been documented. While most changes in HLM reset to basal level after stress, some are inherited through meiosis or mitosis and carry ‘stress memory’ (Luna *et al*., 2012; Slaughter *et al*., 2012). Besides these corollary data, genetic studies implicate HLMTs in plant responses to biotic and abiotic stress factors. Arabidopsis trithorax-like factor ATX1 mediates H3K4me3 and dehydration responses (Ding *et al*., 2011). The *atx1* mutant shows decreased germination rates, larger stomatal apertures, rapid transpiration, and decreased tolerance to dehydration stress. By contrast, Arabidopsis *atx4* and *atx5* mutants that are reduced in H3K4me3 are tolerant to drought stress (Liu *et al*., 2018). Arabidopsis SDG8 is a H3K36 and H3K4 methyltransferase implicated in regulation of flowering time (Zhao *et al*., 2005), shoot branching (Dong *et al*., 2008), carotenoid biosynthesis (Cazzonelli *et al*., 2009), responses to brassinosteroids (Wang *et al*., 2014), regulation of carbon responsive gene expression (Li *et al*., 2015), and RNA processing (Li *et al*., 2020). SDG8 mediated H3K36 methylation is required for defense gene expression (Berr *et al*., 2010), and chromatin remodeling at resistance gene loci (Palma *et al*., 2010). Arabidopsis SDG25, together with SDG8 regulate expression of genes that broadly contribute to plant defense responses (Lee *et al*., 2016). Studies in Arabidopsis have advanced our knowledge on the functions of plant HLMTs but such studies are lacking in crop plant species.

The regulatory and biological functions of tomato HLMTs, SDG33 and SDG34, in biotic and abiotic stress responses were studied. Single and/or double mutants of tomato *SDG33* and *SDG34* genes show increased pathogen resistance and stress tolerance. These responses are underpinned by altered genome wide changes in gene expression, H3K4 and H3K36 trimethylation at chromatin of transcriptional repressors and negative regulators of stress tolerance. In particular, SDG33 and SDG34 are required for pathogen and stress induced enrichment of H3K4me3 and H3K36me3 at target genes that are consistent with both independent and synergistic functions of the two genes. Tomato SDG33 and SDG34 are orthologues of Arabidopsis SDG8. The two tomato genes individually complement the Arabidopsis mutant in SDG8 that shares similar biochemical functions and is an orthologous gene. Intriguingly, mutations in the Arabidopsis and tomato HLMTs display contrasting phenotypes suggesting their distinct biological functions attributed to differences in target gene selection in the two plant genomes. In sum, loss of SDG33 and SDG34 remove the predisposition to stress and pathogen infection, primarily by disrupting the permissive transcriptional context and reducing expression of negative regulatory factors, resulting in elevated pathogen and stress tolerance of the tomato *sdg* mutants.

## Results

### Characterization of *SDG33* and *SDG34* encoding histone methyltransferases

SDG33 (*Solyc04g057880*) and SDG34 (*Solyc06g059960*) encode class II <u>S</u>ET <u>D</u>omain <u>G</u>roup (SDG) HLMTs (Cigliano *et al*., 2013) (Fig. 1a). Similar to other class II SDG proteins, SDG33 and SDG34 carry the evolutionarily conserved SET domain (Pfam ID: PF00856) responsible for their catalytic activity (Zhang & Bruice, 2008). In addition, SDG33 and SDG34 proteins contain an N-terminal <u>A</u>ssociated <u>W</u>ith <u>S</u>ET (AWS) (SMART entry: SM00570), Post SET (SMART entry: SM00508) domains typical of class II HLMTs (Fig. 1a) (Springer *et al*., 2003) and also carry an additional zinc finger domain with conserved cysteine and tryptophan amino acids (zf-CW, PFam 07496). The zf-CW domain binds to DNA and specific histone methylation states (Perry & Zhao, 2003) which suggest that SDG33 and SDG34 function by recognizing and modifying histone methylation patterns. Multiple sequence alignment and phylogenetic analyses reveal that Arabidopsis SDG8 and rice SDG725 are the two class II SDG proteins most closely related to tomato SDG33 and SDG34 (Fig. 1b). SDG33 and SDG34 share only 60.04% amino acid identity when the entire proteins are compared but their SET domains are 100% identical. SDG33 shares 52.91% identity with Arabidopsis SDG8 whereas SDG34 shares 57.11 % identity.

**Fig. 1.**
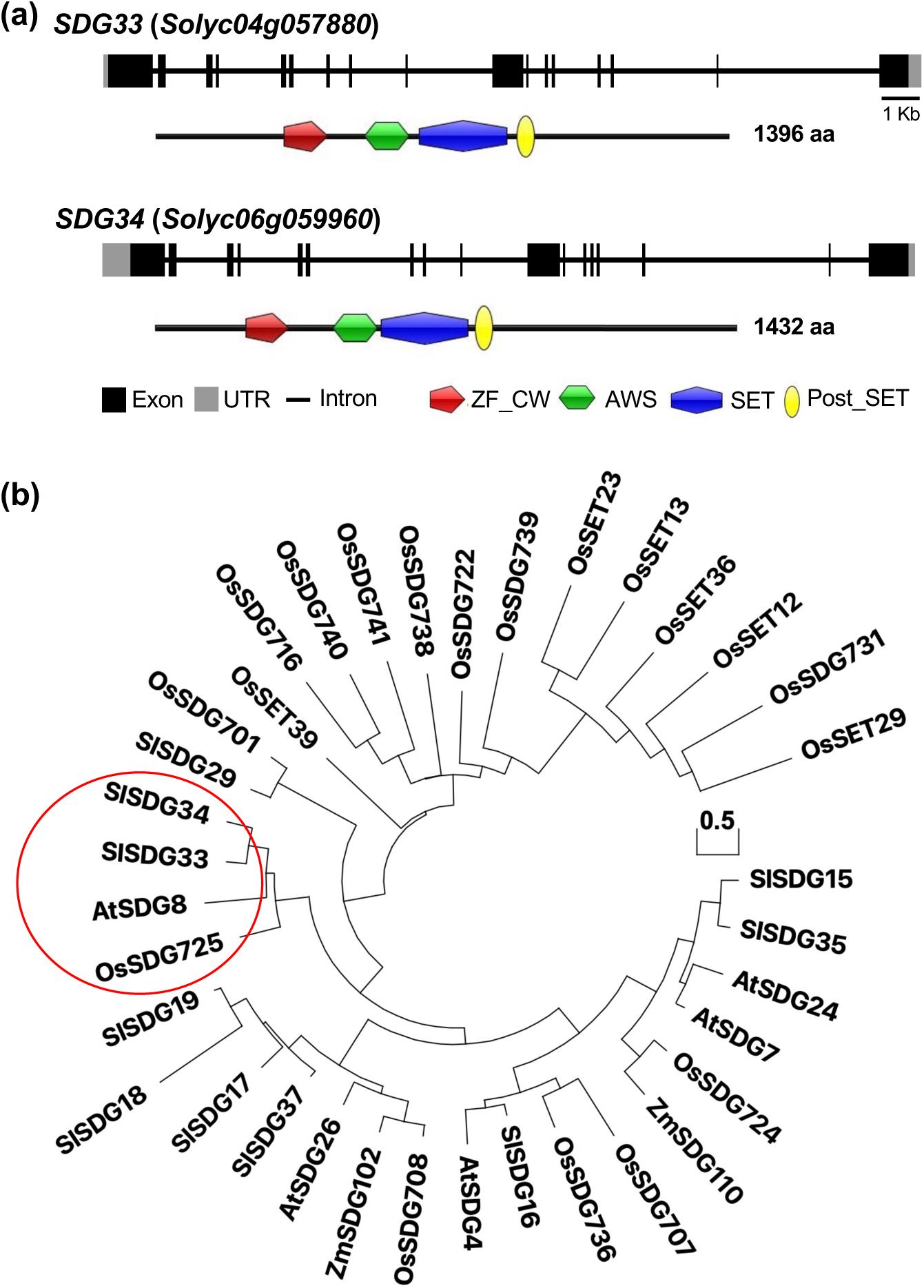
Genomic structures, conserved protein domains and phylogenetic analysis of SDG33 and SDG34. **(a)** Structure of *SDG33* and *SDG34* gene and the encoded proteins showing conserved domains. Black shaded boxes represent exons, the grey shaded boxes represent the UTRs, and introns are shown as horizontal lines. The conserved domains C3HC4 type zinc-finger (ZF-CW) domain, associated with SET domains (AWS), Su(var)3-9, Enhancer-of-zetste, Trithorax (SET) domain, and the Cysteine-rich motif following a subset of SET domains (Post_SET). **(b)** Maximum likelihood phylogenetic analysis of class 11 SDG proteins. Phylogenetic analysis was performed using Mega7 package. The tree is drawn to scale with branch length measured in the number of substitutions per site. Bootstrap values higher than 50% are shown. *Arabidopsis thaliana* (*At*), *Solanum lycopersicum* (*Sl*), *Zea mays* (*Zm*) and *Oryza sativa* (*Os*).

### SDG33 and SDG34 have distinct and overlapping expressions

To gain insight into the biological functions of SDG33 and SDG34, we examined their expression patterns in different tissues using quantitative Reverse Transcription Polymerase Chain Reaction (RT-qPCR) (Fig. 2a). *SDG33* and *SDG34* were both expressed in all tissues tested, with the highest expression in roots and petals, respectively. *SDG33* expression was notably higher in roots followed by petals, which showed 24-fold lower expression compared to roots but about 50-fold higher expression compared to leaves. The expression of *SDG33* in stamens and stigma is higher than in leaves and fruits at all stages of ripening tested. Leaves display the lowest level of *SDG33* expression, while green and red fruit showed about two-fold higher expression than in leaves. *SDG34* expression is notably higher in petals, about 500-fold higher compared to expression in leaves which had the lowest expression of *SDG34*. The stigmas and the stamens exhibit 4-fold higher expression of *SDG34* than leaves.

**Fig. 2.**
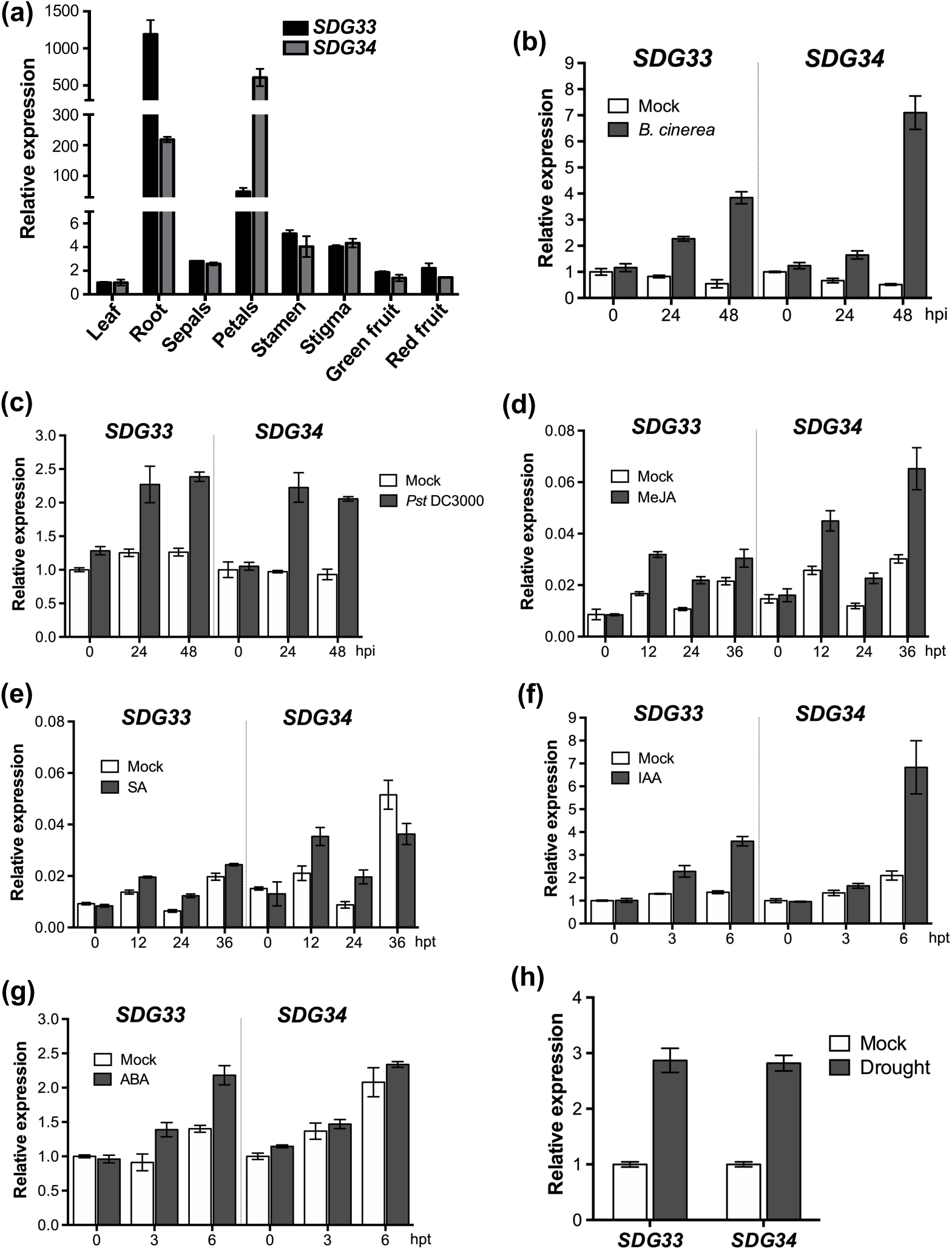
Tissue specific and induced expression of *SDG33* and *SDG34* genes in wild type tomato in response to pathogens, hormones and drought stress. (a) Tissue specific expression of *SDG33* and *SDG34* in wild type tomato plants. Expression of *SDG33* and *SDG34* in response to **(b)** *Botrytis cinerea*, **(c)** *Pseudomonas syringae* pv. *tomato* DC3000 Δ*AvrPto*, **(d)** Methyl jasmonate (MeJA), **(e)** Salicylic acid (SA), **(f)** Indole-3-Acetic Acid (IAA), **(g)** Abscisic acid (ABA), and **(H)** drought stressed plants. Bars represent the means; the error bars represent the standard deviations of three technical replicates of each treatment. The tomato *Actin* gene was used as an internal control in the qRT-PCR experiments. The experiment was repeated at least 2 times with similar results.

In addition, the fungal pathogen *Botrytis cinerea* and the bacterial pathogen *Pseudomonas syringae* induced *SDG33* and *SDG34* genes (Fig. 2b,c). *SDG33* and *SDG34* were highly induced by Methyl-jasmonate (MeJA), slightly induced by salicylic acid (Fig. 2d,e), and indole acetic acid (IAA) (Fig. 2f). *SDG33* is induced by Abscisic acid (ABA) at 3 and 6 h after treatment while *SDG34* expression level was similar to mock treated plants showing some specificity in expression (Fig. 2g). Furthermore, *SDG33* and *SDG34* showed about 3-fold induction after 8 days of drought stress imposed by withholding water (Fig. 2h). Thus, *SDG33* and *SDG34* are transcriptionally regulated by biotic and abiotic stress factors and multiple plant hormones known to mediate plant responses to stress suggesting a broad function. There were some variations in the levels and temporal expression of the two genes but generally both showed the same patterns suggesting overlapping and distinct functions.

### Characterization of tomato *sdg33*, *sdg34* and *sdg33sdg34* mutants

CRISPR/Cas9 mediated gene-edited tomato lines were generated to study the biological functions of *SDG33* and *SDG34* genes. Homozygous lines for two independent mutant alleles of each gene were isolated and characterized (Fig. 3a,b). The *sdg33-64B* mutant allele carried a deletion of 58 bp while *sdg33-64S* had a deletion of 6 bp and a single nucleotide insertion within exon. The deletions in *sdg33-64B* and *sdg33-64S* introduced premature stop codons resulting in truncated proteins. The *sdg34-76* mutant had a deletion of 119 bp and an insertion of 1 bp and *sdg34-27* had a 150 bp deletion and a 4 bp insertion (Fig. 3a,b). All the deletions in *sdg34* mutants were in the last exons and introduced premature stop codons. The expression of *SDG33* and *SDG34* genes were severely reduced in *sdg33* and *sdg34* mutants compared to the WT (Fig. S1a,b) but the transcripts likely produced truncated and non-functional proteins. The expression of *SDG34* in *sdg33* mutant and *SDG33* in *sdg34* was not affected suggesting their expression is independent of each other (Fig. S1c,d).

**Fig. 3.**
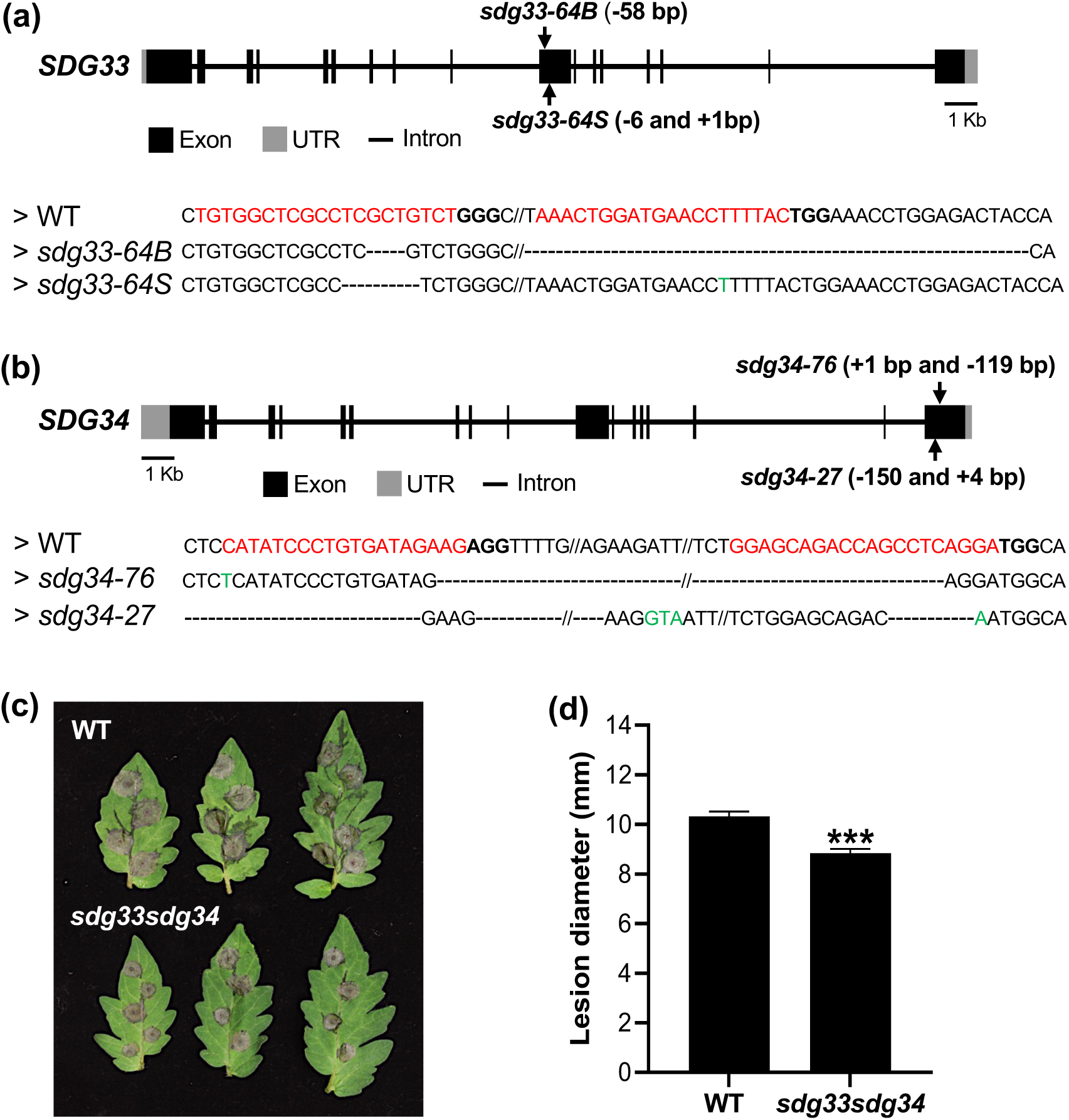
Characterization of tomato *sdg33* and *sdg34* mutants. **(a,b)** Schematics showing the genomic structure of *SDG33* and *SDG34* CRISPR/Cas9 edited mutant alleles. Black shaded boxes represent exons, the grey shaded boxes represent the UTRs and introns are shown as horizontal lines. Alignment of sequences of mutated alleles and the wild type sequences. The mutated alleles include deletions (shown by dashed lines) and insertions (shown by green letters). Only aligned sequences and the mutations are shown. The targets are shown by letters in red and the protospacer adjacent motif (PAM) is shown by the bold-faced letters after the targets. **(c)** Disease symptoms, **(d)** Disease lesion size in wild type and *sdg33sdg34* double mutant. The data shown are the mean ± SE (*n* >30). Different letters represent significant differences among genotypes. Bars represent the means; the error bars represent the standard deviations of three technical replicates of each treatment. The tomato *Actin* gene was used as an internal control in the qRT-PCR. The experiment was repeated at least two times with similar results.

To determine whether SDG33 and SDG34 function redundantly or have distinct functions, *sdg33sdg34* double mutant was generated by crossing the *sdg33-64B* and *sdg34-*76 single mutants. The number and weight of fruits were comparable to wild type plants, but fruits of the *sdg* mutants had a pointy tip at the bottom giving them an oxy heart shape in contrast to the round bottom of the WT fruits (Fig. S1e).

### The distinct and shared functions of orthologues *SET DOMAIN GROUP GENES* from Arabidopsis and tomato

Plants were tested for responses to *B. cinerea* to determine the defense functions of the two tomato genes. The single mutants showed a WT response to *B. cinerea* (Fig. S2) whereas the *sdg33sdg34* double mutant showed increased resistance with significantly smaller disease lesions than the WT (Fig. 3c,d) that suggested functional redundancy between SDG33 and SDG34. Tomato SDG33 and SDG34 are orthologues of Arabidopsis SDG8 that carries similar domain architecture indicative of similar functions (Fig. 1a). The resistance of the tomato *sdg33sdg34* double mutant to *B. cinerea* was intriguing considering the positive role of SDG8 in plant immunity (Lee *et al*., 2016). To delineate these disparities, we generated Arabidopsis transgenic lines that express the tomato genes generating *Sl*SDG33-HA:*sdg8-2* and *Sl*SDG34-HA:*sdg8-2* lines. The Arabidopsis transgenic lines were completely rescued for the smaller plant stature and early flowering phenotypes of Arabidopsis *sdg8-2* mutant (Fig. 4a) (Kim *et al*., 2005). The *B. cinerea* and *Alternaria brassicicola* susceptibility (Fig. 4b-e), and the diminished global H3K4me3 and H3K36me3 levels in Arabidopsis *sdg8-2* mutant were rescued close to wild type levels or fully rescued (Fig. 4f) (Cazzonelli *et al*., 2009; Lee *et al*., 2016). Further, the expression of Arabidopsis *CAROTENOID ISOMERASE 2* (*CCR2*) gene is strictly dependent on SDG8 (Cazzonelli *et al*., 2009) while the tomato *CCR2* was expressed independent of tomato SDG33 and SDG34 (Fig. S3). Interestingly, the diminished expression of *CCR2* in Arabidopsis *sdg8-2* mutant was partially restored in *SlSDG33-HA*:*sdg8-2* and *SlSDG34-HA*:*sdg8-2* Arabidopsis lines with the highest expression of SDG33 restoring *CCR2* expression to almost wild type levels (Fig. 4g). Thus, *SlSDG33* and *SlSDG34* can substitute the HLMs, downstream gene expression, and biological functions of *At*SDG8 suggesting they share biological and biochemical functions.

**Fig. 4.**
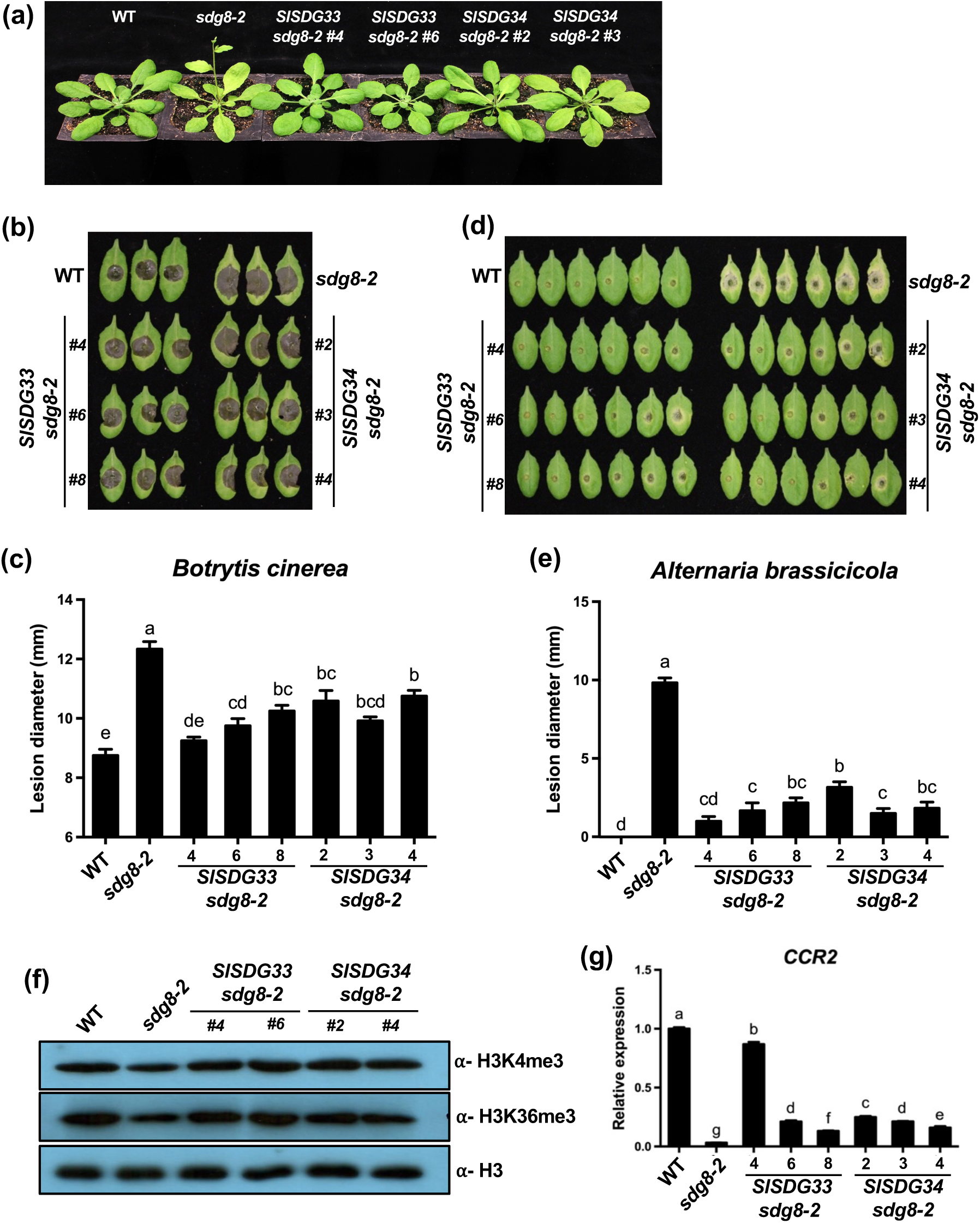
Tomato SDG33 and SDG34 rescue Arabidopsis *sdg8* mutant phenotypes. **(a)** Growth phenotypes of Arabidopsis wild type, *sdg8-2*, and *SlSDG33 or SlSDG34* expressing *sdg8-2* lines (*SlSDG33*;*sdg8-2* and *SlSDG34*;*sdg8-2*). **(b-e)** Fungal resistance of wild type, *sdg8-2*, and *SlSDG33* or *SlSDG34* expressing *sdg8-2* lines. **(b)** Lesion development and **(c)** disease lesion size at 3 days after *Botrytis cinerea* inoculation. **(d)** Lesion development and **(e)** disease lesion size at 4 days after *Alternaria brassicicola* inoculation. Leaves of each line were drop-inoculated with *B. cinerea* (2.5 × 10^5^ spores/mL) or *A. brassicicola* (2.5 × 10^5^ spores/mL) spores. The data shown are the mean ± SE (*n* = 24). Different letters represent significant differences among lines. The statistical significance of differences was analyzed by Tukey’s honestly significant difference (HSD) test (*P* < 0.05). **(F)** The effects of tomato SDG33 and SDG34 methyltransferases on global histone methylation in Arabidopsis transgenic lines. Histone enriched proteins were extracted from 4-week-old plants and global histone methylation levels were determined on Western blot using histone H3 lysine methylation specific antibodies (H3K4me3 and H3K36me3). H3 antibody was used as a loading control. **(g)** *CCR2* gene expression in wild type, *sdg8-2* mutant, SlSDG33 or SlSDG34 expressing Arabidopsis lines (*SlSDG33*;*sdg8-2* and *SlSDG34*;*sdg8-2*) lines. Error bars indicate SD of three technical replicates. The experiment was repeated at least two times, with similar results.

### SDG33 and SDG34 mediate genome-wide transcriptome changes in response to *Botrytis cinerea*

To decipher genome-wide mechanisms underlying the *B. cinerea* resistance of the double mutant, we conducted RNA-seq analysis of *sdg33sdg34* and WT plants using leaf tissues 24 h after mock (control) or *B. cinerea* inoculation. PCA analysis of the transcriptomes indicated a big difference between WT and the *sdg33sdg34* double mutant under the mock condition (Fig. S4a). Therefore, we first analyzed differentially expressed genes (DEGs) regulated by SDG33/SDG34 under the control condition by comparing WT mock and *sdg33sdg34* mock using DESeq2 (Love *et al*., 2014) (adjusted *P*-value < 0.01 and fold change > 2). 165 DEGs were identified, with 98 down-regulated and 67 up-regulated in the double mutant compared to the WT (Fig. 5a,b; Table S1). The 98 down-regulated DEGs formed two clusters: Cluster 1 contains 27 genes that were significantly overrepresented with GO terms related to cellular metabolism such as ‘pigment metabolic processes’ and ‘cofactor metabolic processes’; Cluster 2 contains 71 genes were enriched with GO terms related to biotic stress response such as ‘defense response’ and ‘response to biotic stimulus’ The upregulated genes had no significant GO term (Fig. 5a,b; Table S2).

**Fig. 5.**
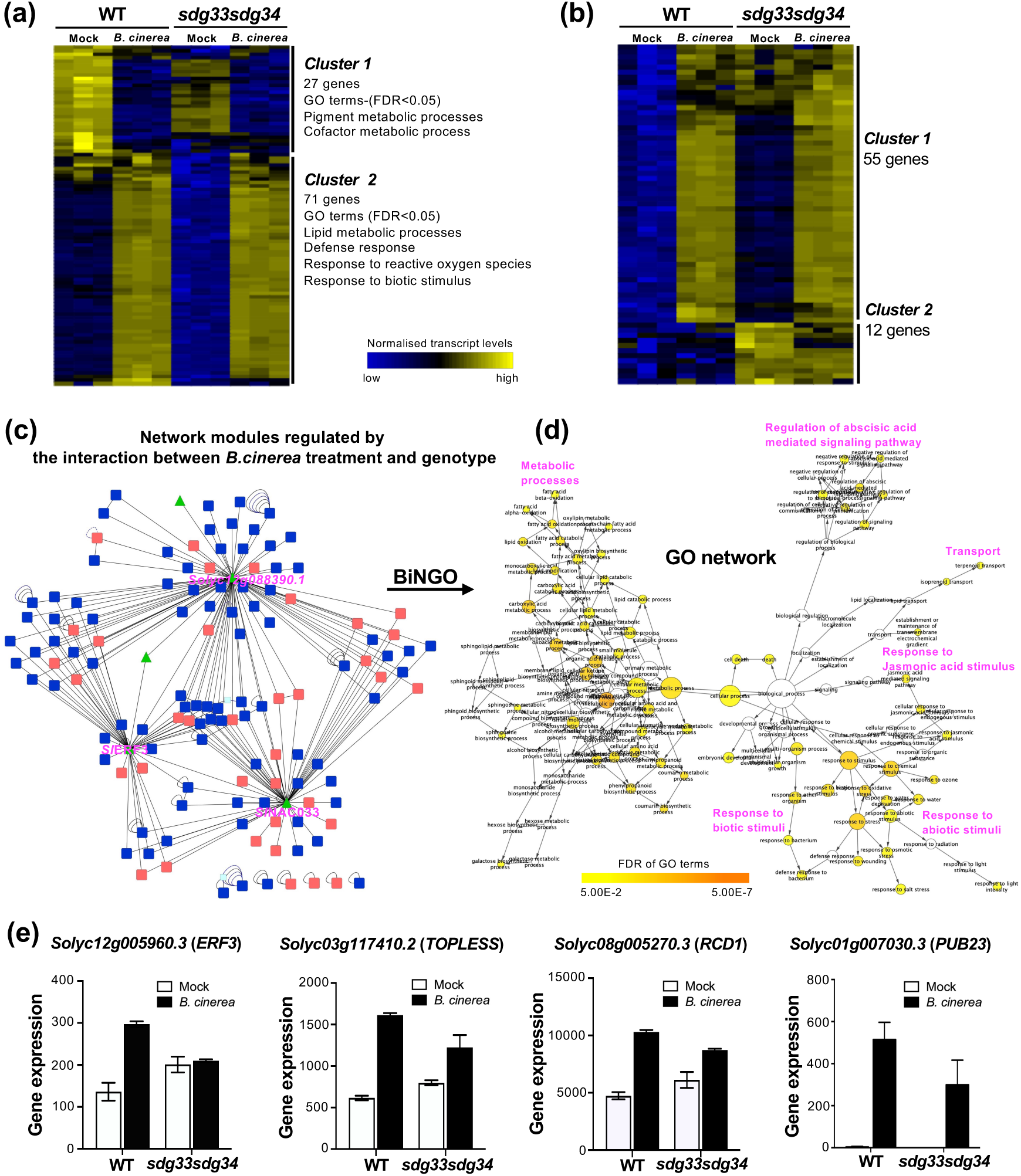
SDG33 and SDG34 regulate genome wide transcriptome in response to *Botrytis cinerea*. **(a,b)** Heatmap showing differential gene expression between *sdg33sdg34* double mutant and wild type plants. **(c)** Gene regulatory network of 179 genes regulated by the interaction between *B. cinerea* and genotype. In this network, the triangle-shaped green nodes show transcription factors (TFs). **(d)** A network view of enriched biological processes from the network generated in **(c)**. Node size is proportional to the number of genes in the test set annotated to the corresponding GO term category. The color of nodes indicates the enrichment *p* value. Orange represents the highest *p* value and yellow represent minimum *p* value above cutoff (FDR < 0.05). White nodes are not significantly overrepresented. **(e)** Expression of transcriptional repressors and other regulatory genes in response to *B. cinerea* infection.

We constructed a gene regulatory network (GRN) from the 165 DEGs using the virtual plant platform (Fig. S4b) (Katari *et al*., 2010). Interestingly, the hub of this GRN is *Solyc09g009490* which is similar to *AT2G36270* that encodes an ABSCISIC ACID-INSENSITIVE 5 (Fig. S4b; Table S3). Recently the barley homolog of ABI5 was shown to mediate drought responses (Collin *et al*., 2020) . In agreement with this, the network was significantly enriched with GO terms related to water stress, such as ‘response to abscisic acid stimulus’ and ‘response to water deprivation’ (Fig. S4c; Table S4). The network was also enriched with GO terms related to defense including ‘immune effector process’ and ‘programmed cell death’ (Padj < 0.05) (Table S4). Indeed, the SDG33/SDG34 regulated DEGs also include *Solyc07g063410* (*SlNAC064*) and *Solyc07g063420.3* (*SlNAC063*) that belonged to the NAC gene family, which have been implicated in diverse functions including biotic and abiotic stress responses (Nuruzzaman *et al*., 2013). Overall, at basal conditions SDG33 and SDG34 regulate a suite of genes that function in metabolism as well as responses to abiotic and biotic stresses.

We next analyzed the RNA-seq data to probe the underlying mechanism of the superior resistance of *sdg33sdg34* double mutant to *B. cinerea*. To this end, we identified DEGs whose *B. cinerea* regulated expression is significantly different between the *sdg33sdg34* mutant and WT, by focusing on the interaction term “genotype: treatment” in model “expression = genotype + treatment + genotype: treatment” using the DESeq2 package (adjusted *p*-value < 0.01). We identified 179 DEGs significantly regulated by the interaction term (Supplemental Table S5) and constructed a GRN from the interaction DEGs (Katari *et al*., 2010) (Fig. 5c; Tables S5, S6). Importantly, three transcription factors were identified as the hubs in this network: (*i*) *Solyc12g088390* encoding a zinc finger protein orthologous to Arabidopsis ZAT10 (AT1G27730), which is a transcriptional repressor involved in abiotic stress responses and JA biosynthesis/signaling (Mittler *et al*., 2006); (*ii*) *Solyc04g009440* (*SlNAC033*), which encodes a NAC domain protein similar to Arabidopsis ANAC002 (AT1G01720, ATAF1), an important regulator of biotic and abiotic stress responses (Jensen *et al*., 2008; Wu *et al*., 2009); and (*iii*) *Solyc12g005960,* which encodes an ETHYLENE RESPONSIVE TRANSCRIPTION FACTOR 3 (*SlERF3/SlERF5.F6*) (Liu *et al*., 2016), orthologous to Arabidopsis *ERF4* (AT3G15210). Arabidopsis *ERF4* is induced by JA and SA (Yang *et al*., 2005), and functions as a transcriptional repressor of JA signaling and resistance to the fungal pathogen *Fusarium oxysporum* (Fig. 5c; Table S6) (McGrath *et al*., 2005). In agreement with this, GO terms “JA mediated signaling pathway” and “response to JA stimulus” were significantly enriched in the interaction network, in addition to other defense-related GO terms such as response to oxidative stress and biotic stimulus (Fig. 5d, Table S7). These GO terms include genes such as tomato *TOPLESS* (*Solyc03g117410*) that is similar to Arabidopsis *TOPLESS*, a known repressor of JA signaling (Pauwels *et al*., 2010), as well as *Solyc08g005270*, an orthologue of Arabidopsis *RCD1* implicated in modulating JA responses (Overmyer *et al*., 2000; Ahlfors *et al*., 2004). In our study, the expression levels of hub gene *SlERF3,* as well as *SlTOPLESS* and *SlRCD1* were induced by *B. cinerea* treatment in the WT, but to a less degree in the *sdg33sdg34* mutant (Fig. 5e). Similarly, the *SlPUB23* gene, encoding a U-box type E3 ubiquitin ligase that likely act as negative regulators of defense (Trujillo *et al*., 2008), was drastically induced by *B. cinerea* in the WT but its upregulation is much lower in the double mutant (Fig. 5e). Overall, our data suggests that attenuated expression of repressors might contribute to the enhanced *B. cinerea* resistance of s*dg33sdg34* mutant. In addition, the interaction genes network was enriched with GO terms related to abiotic stress (Fig. 5c,d; Table S7), again indicating that SDG33 and SDG34 regulate both biotic and abiotic stress responsive genes, potentially mediating crosstalk between different stress responses.

### SDG33 and SDG34 are required for histone H3 lysine 4 and lysine 36 methylation

To directly test the role of SDG33 and SDG34 in HLM in tomato, we analyzed the global H3K4 and H3K36 methylation levels on an immunoblot using antibodies specific to various histone H3 lysine methylations (H3Kme) (Fig. 6a,c). Band intensities were quantified with ImageJ software (Schneider *et al*., 2012) using H3 bands as a reference (Fig. 6b,d). H3K4 and H3K36 mono-methylations were greatly elevated in the *sdg33* mutant while the *sdg34* single and *sdg33sdg34* double mutant was reduced suggesting contrasting impacts on this modification. SDG34 and SDG33 both affect global H3K4 and H3K36 di- and trimethylations with reduced levels in single and double mutants (Fig. 6). In sum, SDG33 and SDG34 have overlapping and distinct functions in the control of H3K4 and H3K36 methylation levels.

**Fig. 6.**
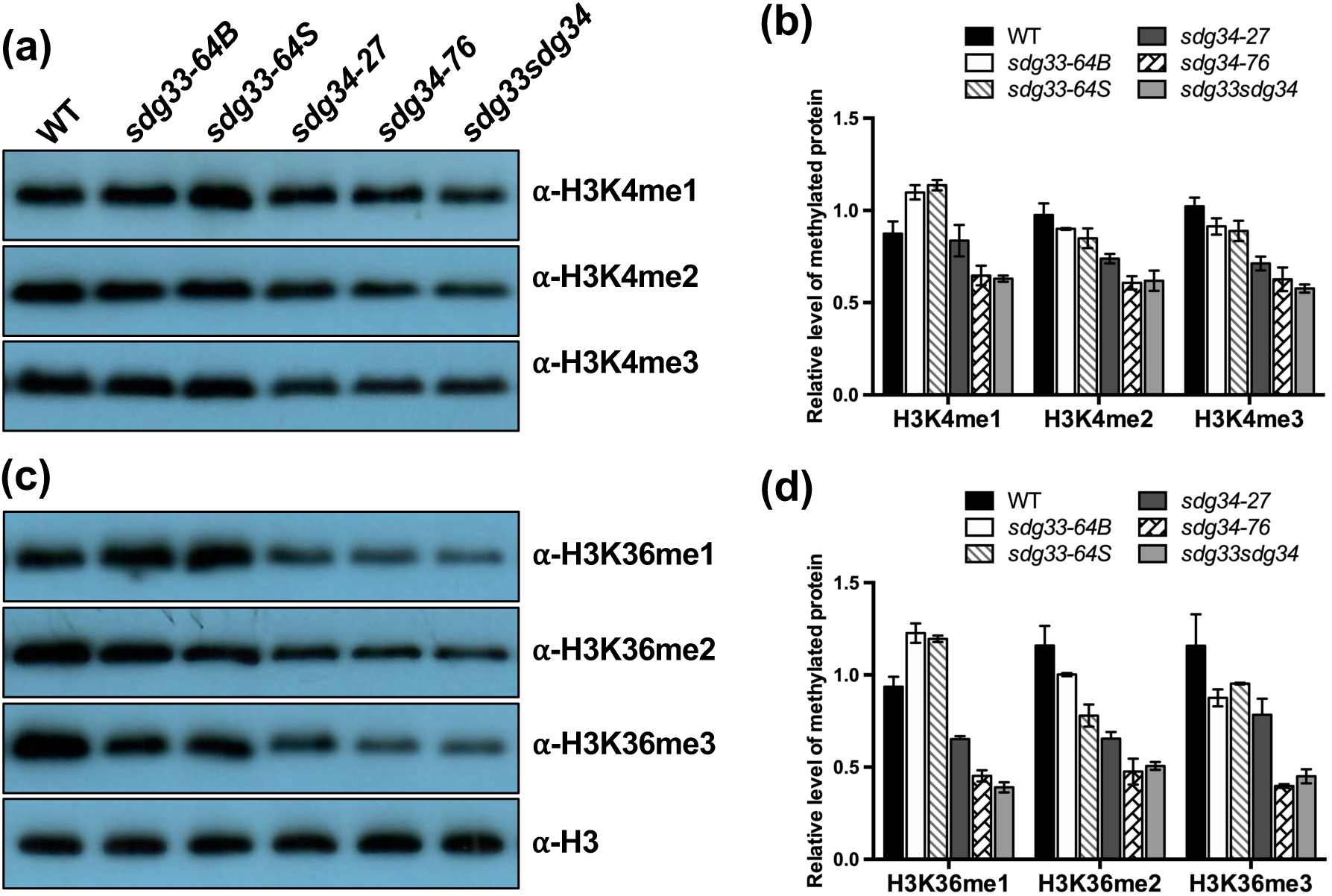
SDG33 and SDG34 regulate global histone lysine methylation profiles in tomato. **(a-d)** Global histone H3 lysine 4 and lysine 36 methylations in tomato wild type, *sdg33, sdg34* single and double mutants. **(b)** and **(d)** quantification of H3K4 and H3K36 methylation levels correspondiong to the Wetsern blots in **(a)** and **(c)**. Bars are means ± SD from two biological replicates. Histone-enriched protein extracts obtained from 4-week-old plants were immunoblotted and probed with antibodies that recognize specific histone methylations. Total amounts of histone H3 are shown as loading controls. Band were quantified using ImageJ software and normalized to H3. Similar results were obtained in two independent experiments.

The transcriptome analyses uncovered genes whose expressions were dependent on *SDG33* and/or *SDG34* (Fig. 5e). To identify changes in methylation status of some of these genes in the *sdgsdg34* double mutant, ChIP experiments were conducted using H3K4me3 and H3K36me3 specific antibodies followed by qPCR with primers at the promoter and exon of the selected genes (Fig. 7a). Rabbit immunoglobulin G (IgG) served as a negative control for immunoprecipitation of all chromatin samples that showed a very low background (Fig. 7b-e). In mock controls, H3K36me3 enrichment at the promoter of *SlERF3*, *SlRCD1* and *SlPUB23*, and H3K4me3 at promoters of *SlTOPLESS* were comparable between WT and *sdg33sdg34* (Fig. 7b-e). The H3K4me3 levels at *SlERF3* promoter and *SlTOPLESS* exon, and H3K36me3 levels at *SlTOPLESS* promoter and exon, and *SlPUB23* exon were lower in the double mutant than WT in control plants, suggesting that these modifications are dependent on SDG33 and/or SDG34 (Fig. 7b-e).

**Fig. 7.**
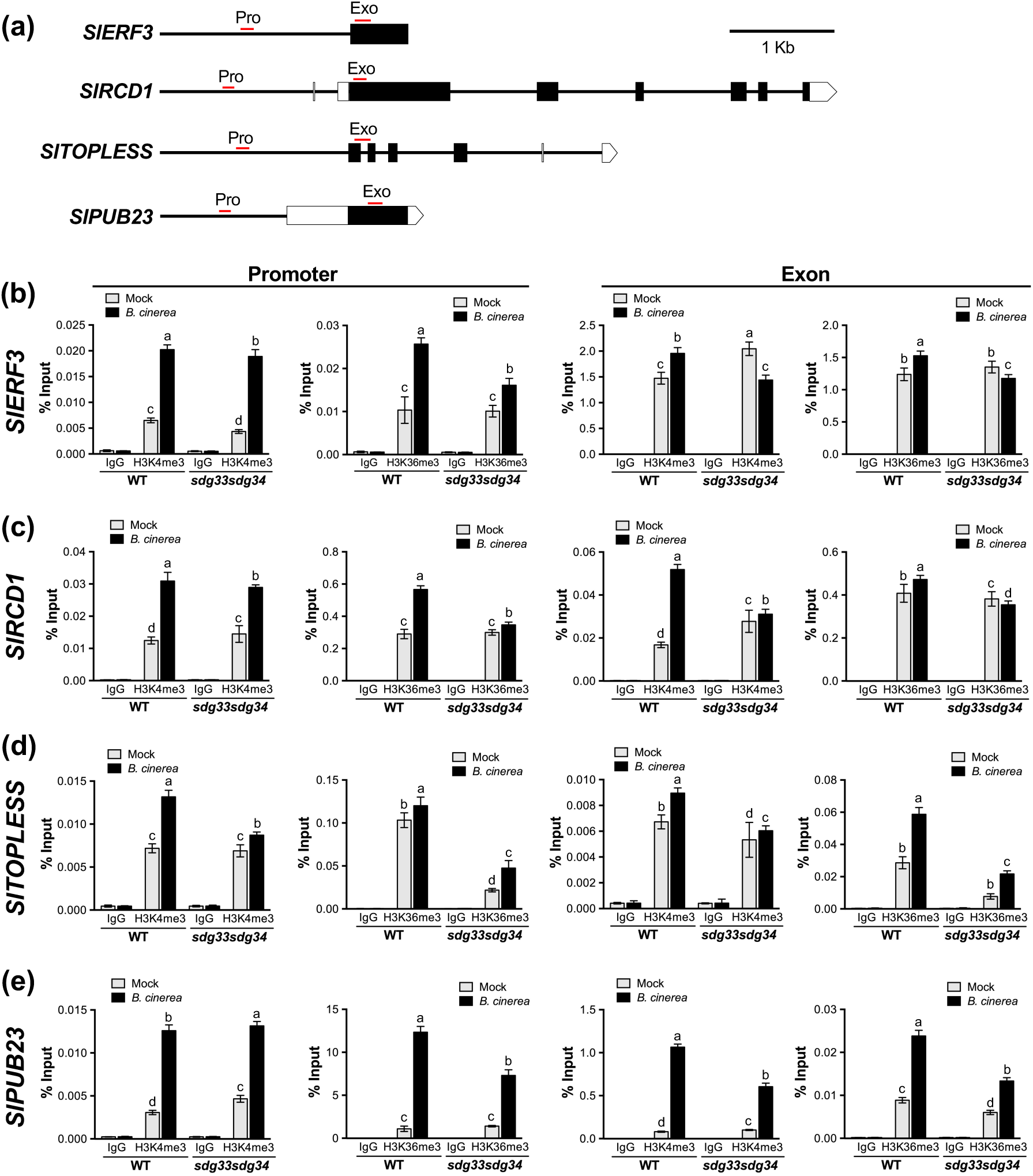
SDG33 and SDG34 regulate the H3K4me3 and H3K36me3 enrichments at target genes in response to fungal infection. **(a)** Schematics showing *SlERF3*, *SlTOPLESS*, *SlRCD1*, and *SlPUB23* genomic region. The position of primers at the promoter (Pro) and coding regions (Exo) used for ChIP-qPCR assays are indicated by the red lines. The solid lines indicate promoter and intron regions, and black boxes indicate exons. **(b-e)** Relative enrichment of H3K4me3 and H3K36me3 at target loci in mock or *B. cinerea* inoculated plants. ChIP was performed on chromatin extracts using antibodies that recognize histone methylations as indicated, and IgG serves as a background level. Data from each experiment were normalized to *ACTIN2* and presented as a percentage of input (% Input). Data are representative of one biological experiment with three technical replicates. Error bars show ± SD (*n* = 3). Different letters indicate significant differences among the genotypes chipped with H3K4me3 or H3K36me3 based on Tukey’s multiple comparisons test (*p* < 0.05). Similar results were obtained in two independent biological experiments. Ab, antibody; WT, wild type.

In response to *B. cinerea*, a significant enrichment of H3K36me3 levels were observed at all the tested loci in WT that was significantly lower in *sdg33sdg34* (Fig. 7b-e). Increased H3K36me3 levels upon *B. cinerea* inoculation at these loci was correlated to their *B. cinerea* induced gene expression in the WT and *sdg33sdg34* mutant (Fig. 5e). Interestingly, both the basal and *B*. *cinerea* induced enrichment of H3K36me3 at promoter and exon of *SlTOPLESS* was significantly lower in *sdg33sdg34* mutant (Fig. 7d). Taken together, SDG33 and SDG34 regulate transcriptional induction by *B. cinerea* through H3K36me3 at *SlTOPLESS*, *SlERF3*, *SlPUB23*, and *SlRCD1* loci.

H3K4me3 levels increased at the promoters and exons of all the genes tested in response to *B. cinerea* in both the WT and the double mutant, except at the *SlERF3* exon, which was decreased by *B. cinerea* (Fig. 7b-e). The increase of H3K4me3 was unexpectedly higher at *SlPUB23* promoters in the *sdg33sdg34* mutant compared to WT. On the contrary, H3K4me3 enrichment at the *SlERF3*, *SlRCD1* and *SlTOPLESS* exon and all the promoter regions were lower in the *sdg33sdg34* mutant than the WT in response to *B. cinerea* (Fig. 7b-e). This enrichment at the promoter and exon regions was correlated to *B. cinerea* induced gene expression (Fig. 5e). Thus, our results indicated that SDG33 and SDG34 impact expression of *SlERF3*, *SlRCD1*, *SlTOPLESS*, and *SlPUB23* genes directly through H3K4me3. Overall, we observed a complex interplay of H3K4me3 enrichment on gene body with a possible role for other HLMTs in regulating H3K4me3 enrichment at specific loci in response to *B. cinerea*.

### Negative regulation of tomato drought stress responses by *SDG33* and *SDG34*

Drought highly induced *SDG33* and *SDG34* genes (Fig. 2h), and SDG33 and SDG34 regulated transcriptome were enriched with drought related genes and pathways suggesting potential roles in abiotic stress responses. To test this possibility, we first assayed tomato seedlings for mannitol induced osmotic stress tolerance (Fig. 8a-e). Mannitol is an osmoticum which lowers water potential of a medium and induces osmotic stress (Claeys *et al*., 2014). On media without mannitol, there were no significant differences in hypocotyl and root length between the mutant and WT seedlings. The *sdg34-76* single and *sdg33sdg34* double mutants showed a significantly higher seedling fresh weight while the other mutants were comparable to the WT (Fig. 8a,c-e). Mannitol significantly reduced seedling growth in both the WT and the mutants but the *sdg33*, *sdg34* single and the *sdg33sdg34* double mutants had significantly longer hypocotyl, root length and higher fresh weight than the WT indicative of osmotic stress tolerance (Fig. 8b-e). The growth of *sdg33sdg34* double mutant seedling on mannitol was comparable to the *sdg33* and *sdg34* single mutants suggesting that SDG33 and SDG34 act through similar mechanisms in regulating osmotic stress. The data show that SDG33 and SDG34 negatively regulate osmotic stress tolerance in tomato seedlings.

**Fig. 8.**
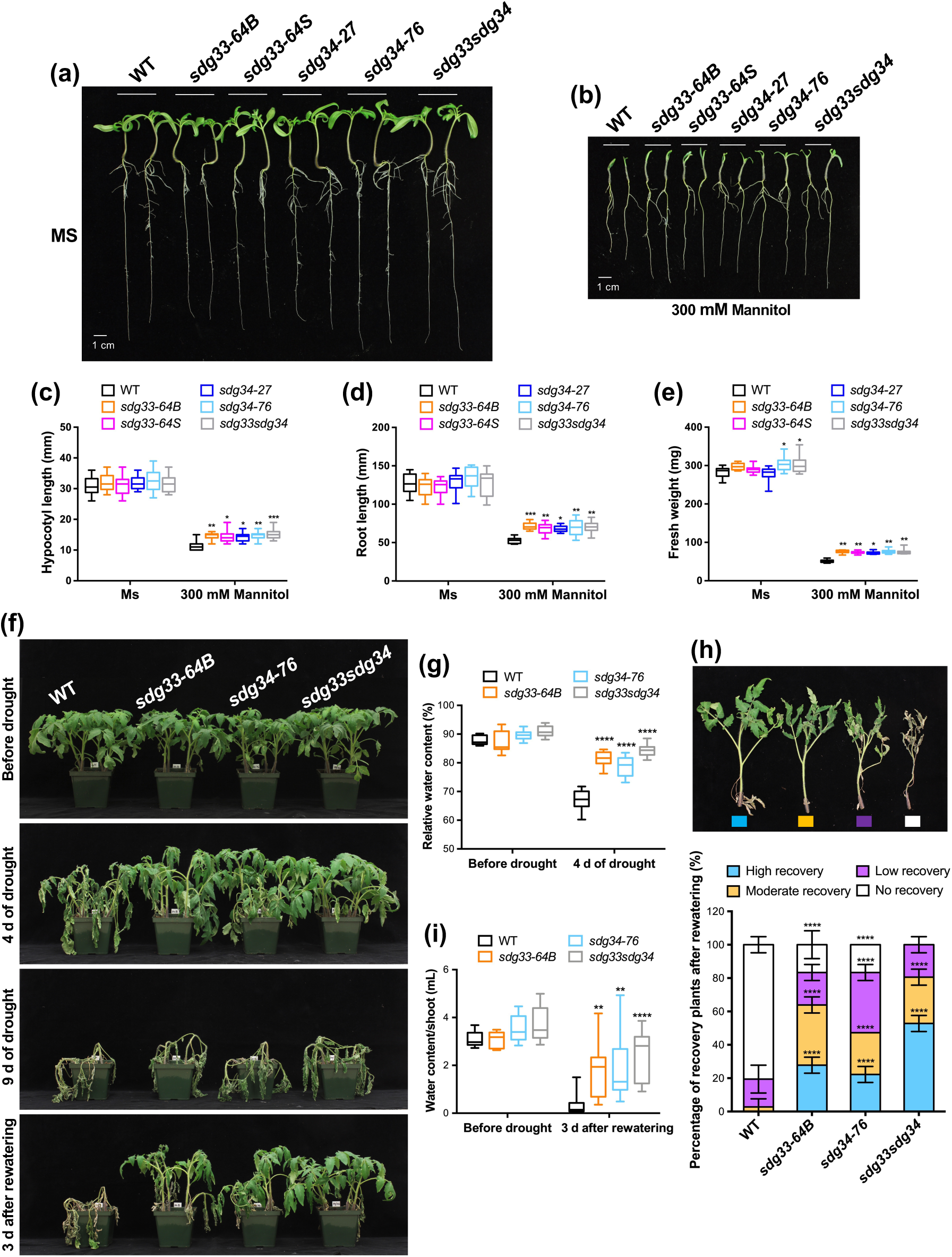
Tomato *sdg33* and *sdg34* mutants show enhanced resistance to osmotic and drought stress. **(a,b)** Seedling growth on media with and without mannitol. Three-day-old seedlings were transferred to MS plate with or without mannitol. **(c)** Hypocotyl length, **(d)** primary root length, and **(e)** seedling fresh weight. Pictures and measurements were taken at 10 days on MS with or without mannitol. The data represent mean ± SE (*n* = 14), the experiment was repeated two times with similar results. Asterisks indicate significant differences (\**p* < 0.05, \*\**p* < 0.01 and \*\*\**p* < 0.001) between the WT and the corresponding mutant line at the same treatment conditions based on Tukey’s multiple comparisons test. **(f)** Growth and extent of recovery after drought stress. Four-week-old plants were subjected to drought stress treatment. **(g)** Relative Water Content at 4 days after drought stress. The data shown are the mean ± SE (*n* = 10). **(h)** Survival rates after re-watering of drought stressed plants. The data shown are mean ± SE (*n* = 36). Data are representative of at least 3 independent experiments. Asterisk indicates significant differences (*p* < 0.0001) between the WT and each genotype according to Tukey’s multiple comparisons test. **(i)** Shoot water content measured at 3 days after re-watering. The data shown are the mean ± SE (*n* = 36). Asterisks indicate significant differences (\*\**p* < 0.01 and \*\*\*\**p* < 0.0001) between the WT and the corresponding mutant line at the same treatment conditions based on Tukey’s multiple comparisons test.

We next investigated the role of SDG33 and SDG34 in drought stress tolerance. Four-week-old soil-grown plants were exposed to drought stress by withholding water for 9 days. After 4 days of drought stress, the WT showed severe wilting symptoms while the single and double mutants showed only slight wilting symptoms with most of the leaves maintaining their turgor (Fig. 8f). Wilting reflects the turgor pressure in a cell, which is highly dependent on the water content of cells (Brodribb & Cochard, 2009). RWC was comparable between mutant and WT plants grown without drought stress. However, in drought stressed plants, the mutants showed significantly higher RWC than the WT with the double mutant having higher RWC than the single mutants (Fig. 8g). The higher water status of *sdg33* and *sdg34* mutants at day 4 corroborated the wilting symptoms (Fig. 8f,g). One possible explanation for the higher water status of the plants could be that stomata closure was greater in *sdg33* and *sdg34* mutants during water stress thereby restricting water loss through transpiration. However, stomata conductance was similar in WT, *sdg33* and *sdg34* mutants throughout the water stress regime. Further, transpiration rate did not differ between the mutants and the WT during stress (data not shown).

Plants were re-watered after 9 days of drought stress and their recovery scored. Drought stressed plants were grouped into different recovery and drought damage severity categories (Fig. 8h). For the WT, 80.5% of plants failed to recover and only 19.5% showed moderate to low recovery. The *sdg33* mutant showed 27.7% high, 36.1% moderate, 19.5% low and 16.7% no recovery categories. The drought stressed *sdg34* mutants were categorized into 22.2% high, 25% moderate, 36.1% low, and 16.7% no recovery. All of the double mutant plants recovered from the drought stress with 52.8% of the plants in the high and 27.8% in the moderate, and 19.4% in low recovery categories (Fig. 8h). Data on drought tolerance of the second mutant alleles of *sdg33* and *sdg34* is presented in Fig. S5. Overall, the single mutants were tolerant to drought, and the double mutant showed significantly higher recovery rate than the single mutants. The higher recovery of the double mutants than the single mutants is consistent with the higher RWC in the mutants during water stress. In the control plants, there were no significant difference between the mutants and the WT in shoot water content but after drought stress, the mutants had significantly higher shoot water content than the WT (Fig. 8i). The improved recovery of the mutants after drought was consistent with the higher shoot water content at the end of drought stress. Taken together, SDG33 and SDG34 act additively to suppress drought stress tolerance.

### SDG33 and SDG34 mediate expression of drought response regulatory genes during PEG induced drought stress

To explore molecular mechanisms of SDG33 and SDG34 mediated suppression of drought tolerance, expressions of potential target genes were studied in response to drought in roots. Different groups of tomato drought responsive genes, mainly encoding transcriptional repressors were identified. This include (1) NAC transcription factors (*SlNAC063* and *SlNAC064*) that are upregulated in the double mutant compared to wild type (Table S1) and were previously implicated in regulation of drought responses (Nuruzzaman *et al*., 2013); (2) Regulatory hubs in the GRN of misexpressed genes in the double mutant compared to WT (*i.e. SlZAT10*, *SlNAC033* and *SlERF3*) as described in previous section (Fig. 5c); and (3) Genes identified from the GRN analysis of misexpressed genes in the double mutant (*i.e. SlPUB23*, *SlRCD1* and *SlTOPLESS*) that are previously implicated as negative regulators of drought responses. In addition, we selected known tomato drought responsive genes such as *Solyc03g025810* (*SlRD29*), a dehydrin orthologous to Arabidopsis *RD29B*, tomato *DEHYDRATION RESPONSIVE ELEMENT BINDING PROTEINS* (*SlDREB2D*) and tomato peroxidase (*SlPOD*) (Gao *et al*., 2020) considered as positive regulators of drought responses in Arabidopsis.

The expression of the above genes was analyzed in roots exposed to 20% PEG-8000 to simulate drought or treated with a regular watering scheme. The basal expression of *SlRD29* was very low in unstressed roots and there was no difference between the WT and mutant plants (Fig. 9). Without stress, significantly lower expression of *SlNAC063*, *SlPUB23*, *SlDREB2D*, and *SlZAT10* was observed in mutants. Expressions of *SlERF3*, *SlTOPLESS*, *SlNAC033*, and *SlNAC064* were lower in unstressed *sdg34* mutant. The changes in gene expression in *sdg33* mutant in unstressed plants were limited and varied depending on the gene that was also mostly reflected in *sdg33sdg34* double mutant (Fig. 9).

**Fig. 9.**
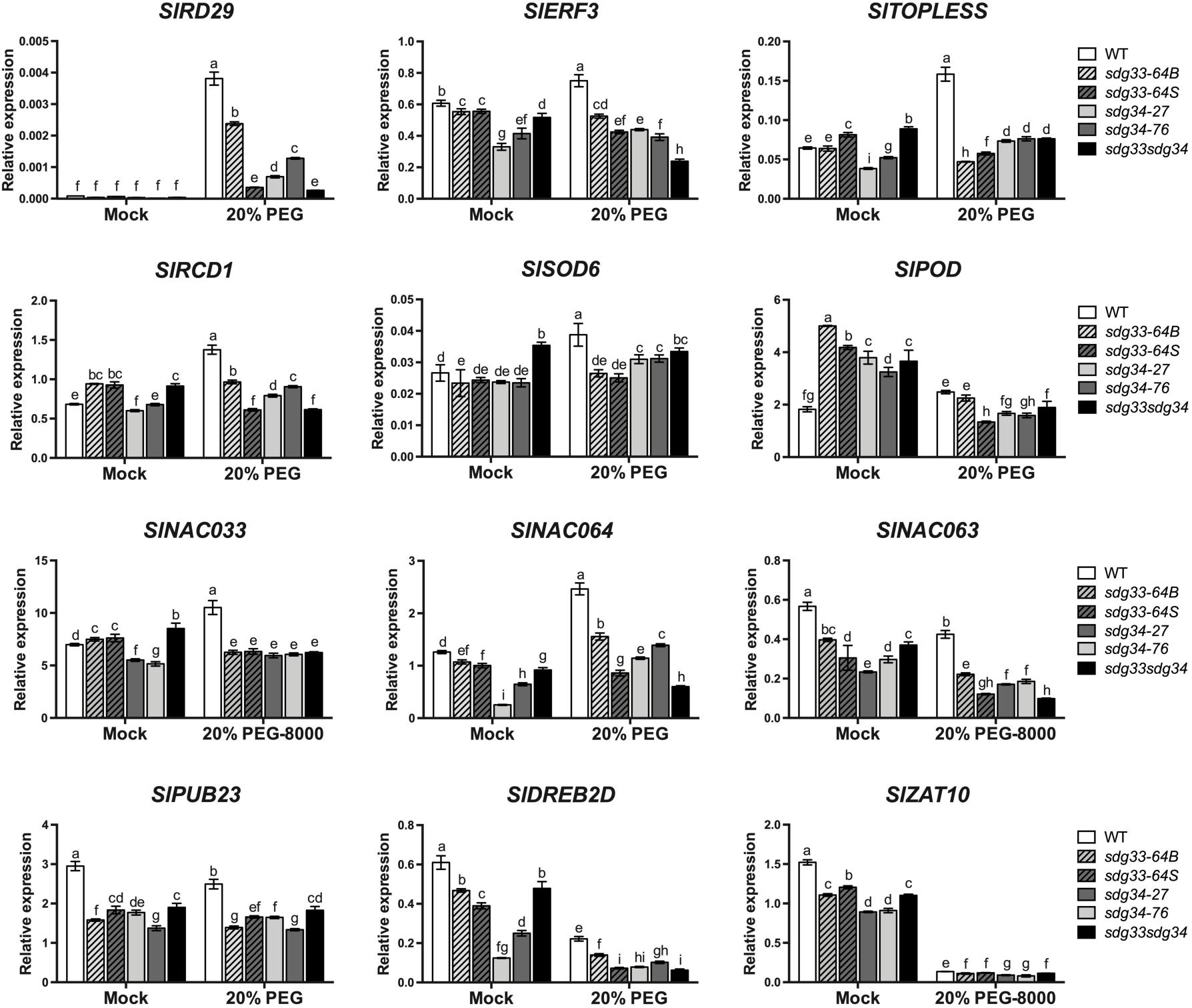
Expression of genes involved in drought stress responses in tomato wild type and mutant lines under drought stress. Expression of *SlRD29*, *SlERF3*, *SlTOPLESS*, *SlRCD1*, *SlSOD6*, *SlPOD*, *SlNAC033*, *SlNAC064*, *SlNAC063*, *SlPUB23*, *SlDREB2D*, and *SlZAT10* in tomato wild type and mutant lines in mock or PEG treated roots. RNAs were extracted from roots of three-week-old wild type and mutant plants after treatemet with water (mock) or 20% PEG-8000. Expression of *SlActin* was used as an internal control. Data represent the means ± SD obtained from experiments performed in triplicates. Different letters indicate significant difference among genotypes (*P* < 0.05; Tukey’s HSD test).

In response to drought, there were two categories of genes based on their expression. The first category includes *SlRD29*, *SlERF3*, *SlTOPLESS*, *SlRCD1*, *SlSOD6*, *SlPOD*, *SlNAC033*, and *SlNAC064* that were significantly induced by drought in WT roots (Fig. 9). *SlRD29* expression was significantly increased by drought in SDG33 and SDG34 dependent manner. While the expression of the other genes in this category was sometimes increased or decreased by drought in the mutants, their expression after drought was generally lower in the mutants than the WT (Fig. 9). The second category of genes includes *SNAC063*, *SlPUB23*, *SlDREB2D*, and *SlZAT10* that were repressed by drought in WT (Fig. 9). Interestingly, *SlNAC063*, *SlDREB2D* and *SlZAT10* were also repressed in response to drought in all the mutants, but their level of expression was significantly lower than the WT (Fig. 9). In *sdg33sdg34* double mutant, *SlTOPLESS*, *SlRCD1* and *SlNAC064* genes were significantly repressed by drought (Fig. 9). The double mutant showed the lowest drought induced expression of *SlRD29*, *SlERF3*, *SlNAC064*, *SlNAC063*, and *SlDREB2D* than the single mutants consistent with synergistic functions of SDG33 and SDG34 in promoting transcription (Fig. 9). Overall, drought-regulated gene expression is dependent on SDG33 and SDG34 consistent with the drought responses of the mutants.

### SDG33 and SDG34 mediate H3K4 and H3K36 trimethylation at drought responsive genes

To understand how SDG33 and SDG34 regulate gene expression, we analyzed H3K4 and H3K36 trimethylation levels at the chromatin of potential target genes (Fig. 10). ChIP-qPCR assay was conducted with H3K4 and H3K36 trimethylation specific antibodies on control and 20% PEG-8000 treated root samples as described in the previous section (Fig. 10). In unstressed plants, the variation in H3K4me3 levels between WT and the mutants was not significant for most genes tested. However, H3K4me3 levels at *SlRCD1*, *SlPUB23* and *SlZAT10* promoter and exons varied significantly between mutants and WT (Fig. 10a). In WT roots, drought significantly enriched H3K4me3 levels at promoters and exons of all the genes tested except at *SlZAT10* where drought depleted H3K4me3. In *sdg33* and *sdg34* single mutants, the H3K4me3 level in response to drought stress at promoters and exons was significantly lower than wild type plants for most genes tested. The *sdg33sdg34* double mutant was significantly reduced in H3K4me3 levels at promoter and exons of most target genes tested relative to the WT. The double mutant showed further reduction relative to the single mutants except at *SlRD29* and *SlERF3* exons and *SlZAT10* promoter (Fig. 10a). SDG33 and SDG34 act independently and additively to modulate H3K4me3 enrichment at target loci.

Similarly, H3K36me3 levels at chromatin of potential target genes were analyzed. In unstressed roots, the H3K36me3 levels were not consistently altered by the *sdg* mutations but the promoters of *SlTOPLESS*, *SlNAC064*, *SlPUB23*, and *SlZAT10* showed lower H3K36me3 enrichment than the WT (Fig. 10b). Reduced levels of H3K36me3 were observed at *SlERF3, SlTOPLESS*, *SlRCD1*, and *SlZAT10* exons in unstressed mutants (Fig. 10b). In PEG treated plants, chromatin at promoters and exons of *SlRD29*, *SlERF3*, *SlTOPLESS*, and *SlNAC064* genes were highly enriched with H3K36me3 in the WT while all the mutant lines showed consistently lower enrichment. The *sdg33* mutant significantly reduced drought induced enrichment of H3K36me3 at promoters and exons of *SlRD29*, *SlERF3*, *SlTOPLESS*, *SlNAC064*, and *SPUB23* (Fig. 10b). In *sdg34* mutant, H3K36me3 was significantly lower than the WT at promoters and exons of *SlRD29*, *SlERF3*, *SlTOPLESS*, *SlNCA064*, *SlPUB23* and promoter of *SlRCD1* suggesting a broader impact than SDG33 (Fig. 10b). H3K36me3 was highly enriched at the promoter and exons of *SlZAT10* compared to the WT chromatin where it was significantly depleted by drought (Fig. 10b). The *sdg33sdg34* mutant showed a generally lower level of H3K36me3 but particularly depleted at *SlRD29*, *SlTOPLESS*, *SlNAC064* promoters and exons, *SlERF3* promoter and *SlPUB23* exon than the single mutant plants (Fig. 10b). There was also differential H3K4me3 and H3K36me3 regulation at *SlRCD1* with no difference observed in H3K36me3 levels but H3K4me3 was enriched in SDG33 and SDG34 dependent manner. In general, SDG33 and SDG34 pose significant regulatory impacts in responses to stress than under normal conditions. There were increases and sometimes decreases in H3K4me3 and H3K36 enrichments after drought in the single and double mutants on target loci. Overall, the trend was a significant reduction of enrichment at most of these loci after drought in the single and double mutants (Fig. 10). The data show that H3K4me3 and H3K36me3 mediated by SDG33 and SDG34 are required for drought inducible gene expression. The repression of drought responsive transcriptional repressors due to loss of the two HLMTs may significantly contribute to the observed drought tolerance of the mutants.

**Fig.10.**
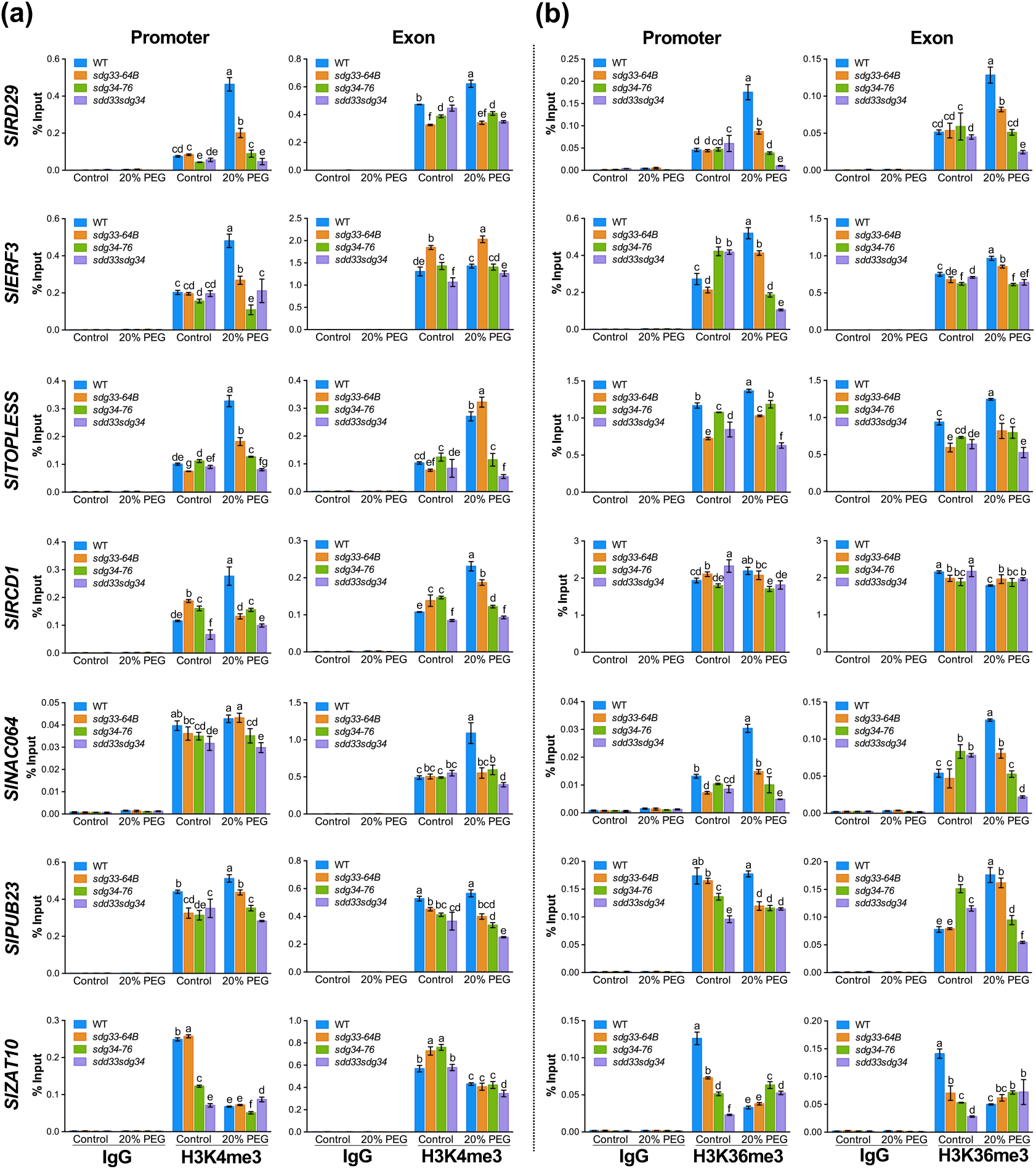
Regulation of H3K4 and H3K36 trimethylation levels at chromatin of SDG33 and SDG34 target genes in response to drought stress. Relative enrichment levels of **(a)** H3K4me3 and **(b)** H3K36me3 chromatin of drought response genes in root tissues. Tomato wild type and mutant plants samples were collected at 3 hours after water (control) or 20% PEG-8000 (PEG) treatments. Nuclei from roots were extracted and ChIP experiments were performed on chromatin extracts using antibodies that recognize histone methylations as indicated, and IgG serves as a background level. Data from each experiment were normalized and are presented as a percentage of input (% Input). Data are representative of one biological experiment with three technical replicates. Error bars show ± SD (*n* = 3). A two-way ANOVA was performed to determine the effects of the tested factors (genotype and treatment) and Tukey HSD test (*p* < 0.05) to compare the means. Different letters indicate significant differences. Similar results were obtained in two independent biological experiments. WT, wild type.

## Discussion

HLMs are major histone epigenetic marks that regulate diverse eukaryotic processes. These modifications are mediated by HLMTs that alter chromatin structure directly or indirectly through reader modules, regulating transcription of genes. Despite this general understanding, the physiological roles of HLMTs and their function in the control of key plant traits through modulation of histone epigenetic marks and the specific enzymes involved are poorly understood. We demonstrate the functions of tomato SDG33 and SDG34 in HLMs, genome wide gene expression and plant responses to biotic and abiotic stress. First, tomato SDG33 and SDG34 redundantly suppress plant immunity. Tomato *sdg33* and *sdg34* mutants show no altered responses to pathogens but the *sdg33sdg34* double mutant exhibited elevated resistance to *B. cinerea*. Second, the *sdg33* and *sdg34* single mutants display enhanced drought stress tolerance, and the double mutant plants were more tolerant than the single mutants. Third, SDG33 and SDG34 affect global H3K4 and H3K36 mono-, di- and trimethylation levels with the *sdg33sdg34* double mutant showing severely attenuated levels. Fungal infection and drought increased H3K4me3 and H3K36me3 enrichment at target genes in SDG33 and SDG34 dependent manner. Additive and independent functions were observed in biochemical and physiological functions of SDG33 and SDG34. Fourth, tomato SDG33 and SDG34 individually substitute the functions of Arabidopsis SDG8. Intriguingly, Arabidopsis *sdg8* and tomato *sdg33* and *sdg34* mutants show distinct phenotypes highlighting the importance of the genomic context and target selection for functions of HLMTs. Reduced enrichment in HLMs in response to drought and fungal inoculation at transcriptional repressor other negative regulatory loci, and attenuation of their expression likely accounts for the observed plant stress tolerance of the tomato mutants highlighting the importance of chromatin profiles for plant stress responses.

This study establishes that SDG33 and SDG34 are tomato HLMTs with largely similar profiles but distinct impacts on H3K4 and H3K36 mono methylations. Also, in the single mutants, the drought induced H3K4me3 and H3K36me3 enrichment was maintained at some of the loci in the single mutants suggesting the presence of other HLMs, and specificity of SDG33 and SDG34 in target selection. Significantly, in the *sdg* mutants, H3K36me3 and H3K4me3 at promoters of target genes were consistently and significantly lower than the wild type tracking their reduced gene expression. Relatively, H3K36me3 was more dependent on SDG33 and SDG34 compared to H3K4me3 suggesting that H3K36 may be the major modification mediated by these SDGs. Regardless, both modifications are active marks of transcription that also accumulate on stress responsive genes (Berr *et al*., 2010; Ding *et al*., 2011; Lee *et al*., 2016). Interestingly, although the biochemical functions of Arabidopsis SDG8 and tomato SDG33 and SDG34 are conserved, the corresponding mutants have contrasting disease resistance functions. Intriguingly, transgenic expression of tomato SDG33 and SDG34 partially or fully rescued the Arabidopsis *sdg8* mutant. These inconsistencies are likely due to differences in key target loci that may be determined by different interactions in tomato and Arabidopsis which partially explain their contrasting impacts on whole plant phenotypes. Arabidopsis SDG8 mediates plant growth and responses to pathogens, at least partially, by regulating H3K4 and H3K36 trimethylation at *CCR2* chromatin and its expression (Cazzonelli *et al*., 2009; Lee *et al*., 2016). Tomato *CCR2* expression was independent of SDG33 and SDG34 suggesting that tomato and Arabidopsis SDG proteins may regulate different target genes.

Plant responses to pathogens and environmental stress entail rapid reprogramming of transcription that occurs in the context of chromatin. The ChIP data suggest that SDG33 and SG34 regulate induced HLM and subsequent reprogramming of gene expression. Network analysis of genes regulated by *B. cinerea* in an SDG33 and SDG34 dependent manner revealed that one of the most regulated genes of this network is *SlERF3,* an ortholog of Arabidopsis *ERF4*. *Sl*ERF3 is a class II ERF protein that has a small Ethylene-Responsive Element Binding Factor associated amphiphilic repression (EAR) motif (DLN xxP) (Liu *et al*., 2016). ERFs containing EAR function to recruit co-repressor proteins such as TOPLESS as well as chromatin remodeling factors to repress target genes (Pan, 2010; Kagale, 2011; Maruyama, 2013). Arabidopsis ERF4 is a negative regulator of JA responsive genes and resistance to the necrotrophic pathogen *Fusarium oxysporum* (McGrath *et al*., 2005). Similarly, Arabidopsis ERF9, an EAR motif protein acts as transcriptional repressor that negatively regulates *B. cinerea* resistance through the repression of ET/JA mediate gene expression (Maruyama, 2013) In Arabidopsis, TOPLESS forms a repressive complex with JAZ proteins that represses the expression JA responsive genes by changing the complex chromatin to a closed one through recruitment of histone deacetylase 6 (McGrath, 2005; Causier, 2012; Ali, 2020). In WT tomato, *SlERF3*, *SlTOPLESS*, and *SlRCD1* are induced by *B. cinerea*, but their induction is lacking or reduced in the *sdg33sdg34* double mutant which correlated with the basal and induced H3K36 methylation at these loci. It is therefore plausible that SDG33 and SDG34 are required to establish the chromatin context necessary to potentiate the induction of *SlERF3* and *SlTOPLESS* in response to *B. cinerea*. Lack of *SlERF3* and *SlTOPLESS* induction is in turn associated with misregulation of JA responsive genes in the *sdg33sdg34* double mutants. This is reflected in the overrepresented GO term, “Response to JA Stimulus” with genes such as *SlRCD1*, in the network of genes regulated by *B. cinerea* in an SDG33 and SDG34 dependent manner. This establishes that SDG33 and SDG34 negatively regulate *B. cinerea* resistance in tomato through H3K4 and H3K36 methylation at chromatin of transcriptional repressors such as *SlERF3* and its co-repressor *SlTOPLESS* whose expression is significantly attenuated in the tomato *sdg* mutants.

SDG33 and SDG34 are major regulators of tomato biotic and abiotic stress responses that act both additively and independently to suppress stress tolerance as evidenced by the better stress tolerance of *sdg33sdg34* double mutant than the single mutants. Interestingly, our RNA-seq data analysis showed that genes regulated by SDG33 and SDG34 under normal conditions as well as in interaction with *B. cinerea* form a network of genes enriched with GO terms related to abiotic stress responses. Analysis of some genes in these networks such as *SlTOPLESS*, *SlRCD1*, *SlERF3*, *SlPUB23*, *SlZAT10*, *SlNAC064*, and *SlNAC063* revealed that they are drought responsive, and their drought induced expression is dependent on SDG33 and SDG34 mediated H3K4me3 and H3K36me3. *Sl*TOPLESS and *Sl*ERF3 are transcriptional repressors (Lin, 2008; Sun, 2008) and Arabidopsis ZAT10, RCD1 and PUB23 negatively regulate osmotic and drought stress responses (Ahlfors, 2004; Mittler, 2006; Cho, 2008). Thus, SDG33 and SDG34 suppress drought tolerance by promoting expression of transcriptional repressors and negative regulators of stress responses. Additional genetic studies are required to validate the functions of these SDG33 and SDG34 target genes. Regulation of drought responsive genes such as *SlRD29*, *SlSOD6*, and *SlDREB2D* demonstrate broader impacts of SDG33 and SDG34 in regulating tomato drought response gene expression. *SlSOD6*, *SlRD29,* and *SlDREB2D* are induced by drought stress but their function in tomato drought stress response is unclear. The contributions of these drought responsive genes, defined based on data from Arabidopsis, may be different or are not sufficient in tomato stress tolerance.

It is noteworthy that while orthology and biochemical functions can be deduced from sequence comparisons between plant species, biological functions may vary greatly. Arabidopsis SDG8, and tomato SDG33 and SDG34 are orthologous with similar biochemical activities but their physiological functions are different. Further studies are required to dissect components that recruit tomato and Arabidopsis HLMTs to target loci and establish unique and conserved components involved in target selection underpinning differences in physiological functions. The observed phenotypes of the tomato mutants are thus likely a result of the decreased expression of critical negative regulators. However, genetic data in tomato is required to establish the contributions of the SDG33 and SDG34 target genes. Interestingly, SDG33 and SDG34 had a greater impact on induced more than basal gene expression, thus induced changes in HLM at target loci serve to integrate responses to environmental cues and potentate plant responses.

## Materials and Methods

### Plant growth, pathogen assays, and molecular analyses

Tomato (*Solanum lycopersicum*) cultivar Castlemart II was the wild-type background of the tomato mutants whereas Arabidopsis *sdg8-2* (SALK_026442) mutant was in the ecotypes Columbia-0. Tomato and Arabidopsis plant growth, pathogen inoculation and hormone treatments, and RNA extraction procedures were described previously (Liao *et al*., 2016; Xu *et al*., 2018). Quantitative-RT-PCR (qRT -PCR) analyses were conducted on iTaq Universal SYBR Green Supermix (Bio-Rad) with the CFX96 qPCR machine (Bio-Rad). Expression levels were calculated by the comparative cycle threshold (Ct) method. Primers used for qRT-PCR are listed in Supporting information Table S8. Tomato SDG33 and SDG34 full length coding sequence (CDS) under their respective native promoter were transformed into Arabidopsis *sdg8-2* (SALK_026442) mutant. Transgenic plants were screened on growth media supplemented with herbicide Basta (Mengiste *et al*., 1997).

### Drought tolerance assays

For osmotic stress tolerance seeds were surface sterilized with 20% Sodium hypochlorite, washed seven times in distilled water and germinated on 1/2 Murashige and Skoog (MS) (PhytoTech Labs) medium with 2% (w/v) sucrose and 0.3% (w/v) Gelzan™ (PlantMedia™) under at 16-hour light, 8-hour dark at 24°C for three days. When the radicle emerged, they were transferred to full MS medium supplemented with 300 mM Mannitol. Seedling length and weight were measured after 10 days. For drought assays on soil, 4 plants per genotype were grown in a square pot and ten pots per genotype with enough water for 4 weeks, drought stress was induced by withholding water for a period of 9 days. Plants were re-watered and survival rate was calculated daily for three days. To measure Relative Water Content (RWC), leaves were collected at day 4 after drought was induced, weighed to get fresh weight (LFW), incubated in water petiole down in a 50 ml conical tube overnight. Leaves were blotted dry and weighed to get saturated weight (LTW). Leaves were then dried at 60°C for seven days and weighed to get dry weight (LDW). RWC was calculated as (LFW-LDW)/(LTW-LDW). Shoot water content was determined for the whole shoot (leaves and stems) according to the formula FW−DW, where FW is the fresh weight and DW is the dry weight obtained oven dried for 3 days at 60°C.

### Global methylation and Chromatin immunoprecipitation (ChIP)-qPCR

Core histone proteins were extracted from 4 weeks old plants and used for Western blot analysis as described previously (Lee *et al*., 2016). The blots were probed with primary antibodies: H3K4me1 (07-436; MilliporeSigma), H3K4me2 (07-030; MilliporeSigma), H3K4me3 (07-473; MilliporeSigma), H3K36me1 (ab9048; Abcam), H3K36me2 (07-369-I; MilliporeSigma), H3K36me3 (ab9050; Abcam), and H3 (ab1791; Abcam) was used as a loading control. Anti-rabbit was used as the secondary antibody. ChIP experiments were performed, as described previously (Lee *et al*., 2016) with minor modifications. For drought-stressed root samples, plants were grown for 3 weeks in Turface Athletics MVP, transferred to 30 ml of 20% PEG-8000 (Polyethylene Glycol) and along with controls (30 ml of water), and treated for 3 h. Roots and leaf tissues (1 to 1.5 g) were cross-linked with 1% (v/v) formaldehyde under vacuum. The tissues were ground to a fine powder in a mortar with liquid nitrogen and chromatin complexes were isolated. Subsequently, the chromatin samples were sonicated using Bioruptor® Pico sonication device (diagenode) to yield fragments between 200-1,000 bp. Precleared chromatin samples by adding Protein A Agarose/Salmon Sperm DNA beads (MilliporeSigma) were incubated with the appropriate antibodies to histone lysine methylation at 4°C overnight and immune complexes were precipitated using Protein A Agarose/Salmon Sperm DNA beads. Immunoprecipitated and input samples were reverse cross-linked at 65°C overnight and samples were digested with 10 μL of 0.5 M EDTA, 20 μL of 1 M Tris-HCl, pH 6.5, and 1 μL of proteinase K (20 mg mL^−1^; Thermo Fisher Scientific) at 45°C for 1 h. Immunoprecipitated DNA samples were purified using PCR purification column (MACHEREY-NAGEL Inc.) and eluted in 80 μL double-deionized water. In qPCR, 2 μL of DNA was amplified using PowerUp™ SYBR™ Green Master Mix (Thermo Fisher Scientific) with specific primers as listed in Table S8. The data is presented as percentage of input values. The antibodies used for the ChIP experiments were: H3K4me3 (07-473, MilliporeSigma), H3K36me3 (ab9050, Abcam) and Rabbit IgG (sc-2027, Santa Cruz) as a negative control.

### Phylogenetic analysis

Tomato protein sequences were obtained from https://solgenomics.net/, *Arabidopsis thaliana (At*) from https://www.arabidopsis.org/, *Zea mays* (*Zm*) sequences were obtained from https://www.maizegdb.org/, and *Oryza sativa* (*Os*) sequence from http://rice.uga.edu/. Maximum Likelihood phylogenetic analysis of class 11 SDG protein from *At*, *Sl*, *Zm,* and *Os* was conducted. These sequences were used to build a phylogenetic tree using the software Molecular Evolutionary Genetics Analysis version 7.0 (MEGA7) (Kumar *et al*., 2016). First, the sequences were aligned using the software MUSCLE with default values. The aligned sequences were used to construct a phylogenetic tree using Maximum Likelihood method and JTT matrix-based model (Jones *et al*., 1992). Initial tree(s) for the heuristic search were obtained automatically by applying Neighbor-Join and BioNJ algorithms to a matrix of pairwise distances estimated using the JTT model, and then selecting the topology with superior log likelihood value.

### Generation of *sdg33* and *sdg34* mutants

To generate *sdg33* and *sdg34* mutants, two specific guide RNA (gRNA) of SDG33 and SDG34 were designed (Table S8) using the CRISPR Plant gRNA design software https://www.genome.arizona.edu/crispr/CRISPRsearch.html. The two gRNA were cloned into the CRISPR/Cas9 vector pKSE401 according to (Xing *et al*., 2014). The vector was electroporated into *Agrobacterium tumefaciens* strain GV3101. Tomato seeds were germinated on germination medium (MS salts, 1X Nitsch’s vitamins, 1.5% sucrose, and 0.25% gelrite). Agrobacterium was grown in YEP medium overnight at 28°C shaking and diluted to OD_600_ = 0.8-1.00 with fresh YEP containing acetosyringone (375 μM). Seven-day-old cotyledons explants were suspended in *Agrobacterium* for 30 minutes and the explants blot dried on sterile blot paper to remove excess *Agrobacterium*. *Agrobacterium* infected explants were co-cultivated on co-cultivation medium (MS salts, 3% sucrose, 1X Nitsch’s vitamins, 0.1 μg kinetin, 200 μg 2.4D, 1 μg 1-Naphthaleneacetic acid (NAA), 1 μg 6-Benzylaminopurine (BA), 200 μM acetosyringone, and 0.25% gelrite) and incubated in the dark at 21°C for 48 hours. After co-cultivation, the explants were transferred to callus induction medium (MS salts, 3% sucrose, 1X Nitsch’s vitamins, 2 μg zeatin, 250 μg timentin, 250 μg cefotaxime, 75 μg kanamycin and 0.25% gelrite) on petri dishes adaxial side up. Explants were transferred into fresh shoot regeneration medium every week. Compact green callus was then transferred to shoot induction medium (MS salts, 3% sucrose, 1X Nitsch’s vitamins, 1 μg zeatin, 250 μg timentin, 250 μg cefotaxime, 75 μg kanamycin and 0.25% gelrite). When shoot had elongated to 2-4 cm, they were excised from callus and transferred to rooting medium (MS salts, 3% sucrose, 1X Nitsch’s vitamins, 1 μg Indole-3-butyric acid (IBA), 250 μg timentin, 75 μg kanamycin, and 0.25% gelrite). Fully rooted plants were transferred into the soil and genotyped by PCR with primers covering the expected deletion. The deletions were confirmed by sequencing. The *sdg33sdg34* double mutants were generated by crossing *sdg33-64B* and *sdg34-76*. The F2 was genotyped by PCR and confirmed by sequencing. Homozygous double mutants were used in the experiments.

### RNA sequencing and data analysis

RNA samples were extracted using the Trizol (Invitrogen) according to the manufacturer’s instructions and submitted to Novogene for RNA-Seq library preparation and sequencing on an Illumina Novaseq platform with paired-end 50 bp format. On average, approximately 43 million read pairs were generated for each library. The raw sequencing reads were trimmed by the fastp software to remove adaptors and low-quality reads (Shifu Chen, 2018). The trimmed reads were then mapped to the latest assembled tomato reference genome SL4.0 using BBmap (Bushnell *et al*., 2017). The overall mapping rate is 97.78% and the multi-mapping rate is about 4.16%. The raw gene counts were generated by FeatureCounts using the tomato gene models ITAG4.0 (Liao *et al*., 2014). Finally, the raw gene counts were processed by DESeq2 (Love *et al*., 2014) to detect the differentially expressed genes (DEGs) with the design (∼ Genotype + Treatment + Genotype:Treatment). DESeq2 also performed an internal gene normalization and low counts filtering for the analysis. The GO analyses for the DEGs were performed using AgriGO database with FDR < 0.05 (Du *et al*., 2010).

To identify possible regulatory relationships and the most regulated genes among DEGs, Arabidopsis Multinetwork database that contains known gene-to-gene interactions in Arabidopsis was used to build the gene regulatory network using the bioinformatics platform VirtualPlant (Katari *et al*., 2010). To do this, the Arabidopsis homologous genes for each tomato gene were identified using BLASTP (E-value cutoff of 1e-06). Next, the Arabidopsis homologs of tomato DEGs were uploaded to the VirtualPlant platform to query the multinetwork database, which includes previously identified protein-protein interactions, transcriptional and post-transcriptional regulation, and TF-gene interactions (DAP-Seq), to construct gene regulatory network (Katari *et al*., 2010). The resulting gene networks were then visualized and analyzed in Cytoscape (Maere *et al*., 2005). To identify the master regulators (hub genes or most regulated genes) in each group, the top 10% of genes with the highest degree in each network were identified as the most regulated gene by using the Network Analyzer function in Cytoscape. To annotate the function of each network, the BiNGO (the Biological Networks Gene Ontology) tool in Cytoscape was used to perform network GO enrichment analysis (Maere *et al*., 2005).

### Statistical analysis

Analysis of Variance (ANOVA) was performed to test the impact of genotype, nitrogen treatment and their interaction on each phenotypic trait. ANOVA was performed in the R software, version 3.3.1 (R Core Team, 2016) using the Agricolae package. Statistical significance was determined at the level (*P* ≤ 0.05).

To determine which means are significantly different, means were compared using the LSD method in the JMP software package. Basis on significance was assessed by least significance differences (*p* = 0.05).

### List of genes and accession numbers

*SlSDG33* (*Solyc04g057880*), *SlSDG34* (*Solyc06g059960*), *SlPUB23* (*Solyc01g007030.3*), *SlRCD1* (*Solyc08g005270.3*), *SlTOPLESS* (*Solyc03g117410.2*), *SlERF3*/*SlERF5.F6* (*Solyc12g005960*), *SlNAC063* (*Solyc07g063410.3*), *SlNAC064* (*Solyc07g063420*), *SlNAC033*, (*Solyc04g009440*), *SlRD29* (*Solyc03g025810*), *SlPOD* (*Solyc10g076240*), *SlSOD6* (*Solyc03g095180*), *SlCCR2*/*CRTISO* (*Solyc10g081650.2*), *AtSDG8* (*AT1G77300*), *AtCCR2*/*CRTISO* (*AT1G06820*).

## Data availability

The RNA-Seq data is deposited in NCBI SRA under accession number PRJNA755600.

## Acknowledgments

This research was funded by grants from NSF (IOS-1916893) and the US-Israel Binational Agricultural Research and Development Fund (BARD) grant IS-5261-20C.

## Author contributions

C.B. and S.L. equally contributed to designing research and conducting the experiments, performing the data analyses, and writing the manuscript. L.T. conducted RNA-seq data analysis. Y.L. wrote the paper, designed the research, and supervised the RNA-seq data analyses. M.V.M. designed and supervised the drought response and physiology experiments. T.M. wrote the paper, developed the project, and designed and supervised research.

## Supporting Information

The following supporting Information are available.

**Fig. S1. Expression of SDG genes and fruit phenotypes in tomato *sdg33* and *sdg34* mutants.**

Expression of **(a)** *SDG33* and **(b)** *SDG34* in tomato WT and mutant plants. Relative expression of **(c)** *SDG33* in *sdg34* mutants and **(d)** *SDG34* in *sdg33* mutants. Bars represent the means; the error bars represent the standard deviations of three technical replicates of each treatment. The tomato Actin gene was used as an internal control in the qRT-PCR experiments. **(e)** Tomato fruits from mutant and WT plants. The experiment was repeated at least two times with similar results and the representative experiment is shown.

**Fig. S2. The single *sdg33* and *sdg34* mutants show no altered pathogen response.**

**(a)** Disease symptoms and **(b)** lesion size on WT, *sdg33* and *sdg34* after inoculation with *Botrytis cinerea* at 3 days after inoculation. The data are the mean ± SE (*n* >36). Leaves of WT and mutant plants were drop-inoculated with *B. cinerea* spores (2.5 × 10^5^ spores/mL). The experiment was repeated three times, with similar results.

**Fig. S3. Tomato *CAROTENOID ISOMERASE2* expression is independent of SDG33 and SDG34.**

Expression of *CCR2* gene in mock- or *Botrytis cinerea* inoculated WT and *sdg33sdg34* based on RNA-seq transcript count data. Error bars indicate the standard deviation of three libraries.

**Fig. S4. Transcriptome analysis in mock and *Botrytis cinerea* inoculated tomato leaf tissues.**

**(a)** Principal component analysis (PCA) of the normalized RNA-seq data; and Gene regulatory network of the 179 genes regulated by genotype. In this network, the triangle-shaped green nodes show transcription factors (TFs). **(c)** A network view of enriched biological processes from the generated network in **(b**).

**Fig. S5. The tomato *sdg33* and *sdg34* single and double mutants are tolerant to drought stress in tomato.**

**(a)** Growth and survival of plants before and after drought stress. Four week-old plants were subjected to drought treatment. **(b)** Relative Water Content at day 4 of water stress. The data shown are the mean ± SE (*n* = 10). **(c)** Survival rates and extent of recovery after re-watering of drought stressed plants. The data shown are the mean ± SE (*n* = 36). Data shown is a representation of experiments repeated at least 3 times. Asterisk indicates significant differences (*p* < 0.0001) between the WT and each genotype according to Tukey’s multiple comparisons test. **(d)** Shoot water content measured at 3 days after re-watering. The data shown are the mean ± SE (*n* = 36). Asterisks indicate significant differences (\**p* < 0.05, \*\**p* < 0.01, \*\*\**p* < 0.001, and \*\*\*\**p* < 0.0001) between the WT and each mutant line counterpart at the same treatment conditions based on Tukey’s multiple comparisons test.

**Table S1. Genes differentially expressed in sdg33sdg34 mock compared to the WT mock.**

**Table S2. Overrepresented GO terms in DEGs downregulated in sdg33sdg34 mock compared to the WT mock.**

**Table S3. Network of downregulated DEGs in sdg33sdg34 compared to the WT.**

**Table S4. Significant GO terms enriched among the downregulated genotype DEGs network.**

**Table S5. Genes whose *Botrytis cinerea* induced regulated expression is significantly different between the WT and the *sdg33sdg34* mutant.**

**Table S6. Network of genes whose *Botrytis cinerea* induced regulated expression is significantly different between the WT and the sdg33sdg34 mutant.**

**Table S7. Significant GO terms enriched in the network of genes whose regulation by the interaction of *Botrytis cinerea* and genotype.**

**Table S8. Primers used in this study.**

