## Supplemental Figure S1-S5 for "Improved pathogen and stress tolerance in tomato mutants of SET domain histone 3 lysine methyltransferases"

**This PDF file includes:**

Fig. S1 to S5

**Other supporting Information for this manuscript include the following:**

Tables S1 to S8

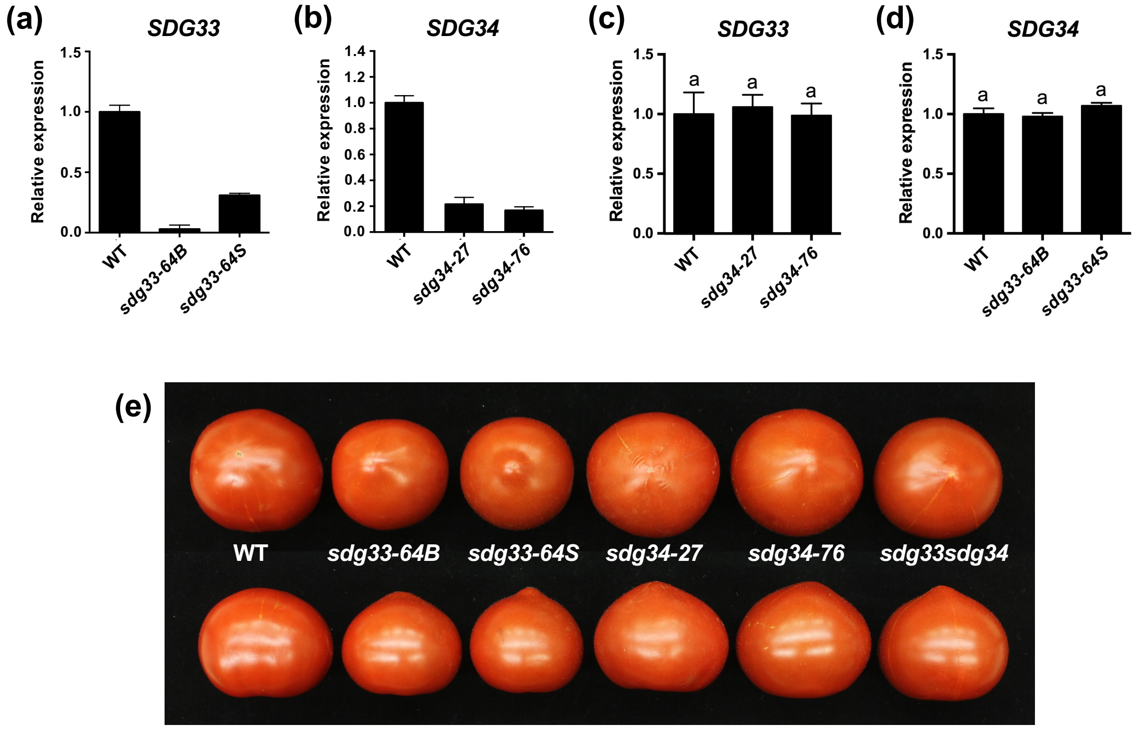


**Fig. S1. Expression of SDG genes and growth phenotypes in tomato *sdg33* and *sdg34* mutants.**
Expression of **(a)** *SDG33* and **(b)** *SDG34* in tomato WT and mutant plants. Relative expression of **(c)** *SDG33* in *sdg34* mutants and **(d)** *SDG34* in *sdg33* mutants. Bars represent the means; the error bars represent the standard deviations of three technical replicates of each treatment. The tomato Actin gene was used as an internal control in the qRT-PCR experiments. **(e)** Tomato fruits from mutant and WT plants. The experiment was repeated at least two times with similar results and the representative experiment is shown.

**(a)**

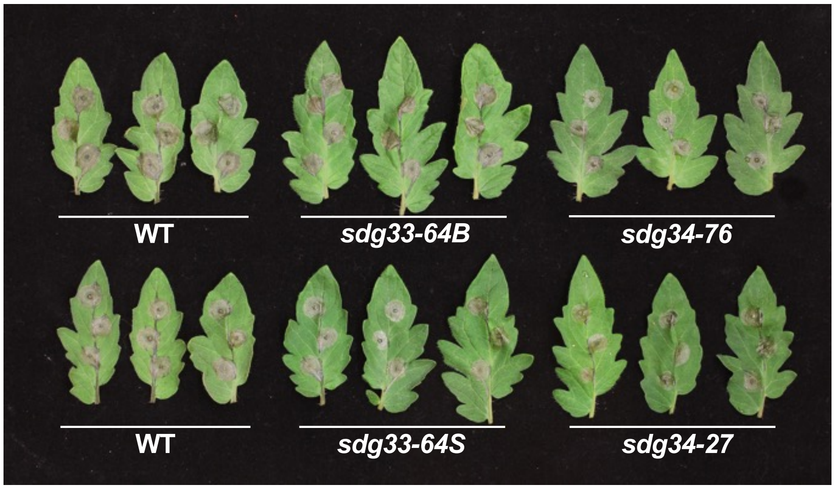


**(b)**

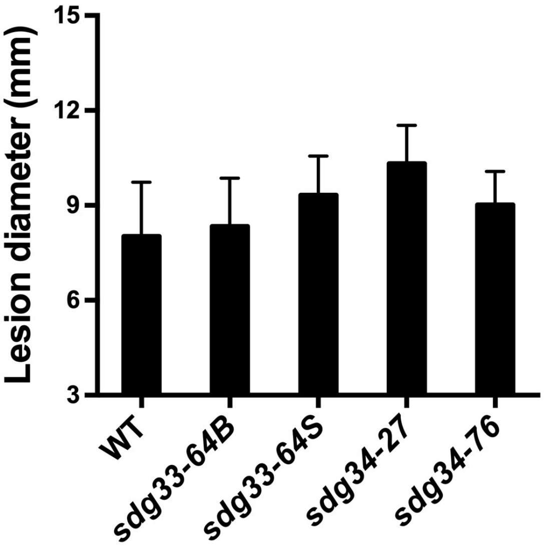


**Fig. S2. The single *sdg33* and *sdg34* mutants show no altered pathogen response.**
**(a)** Disease symptoms and **(b)** lesion size on WT, *sdg33* and *sdg34* after inoculation with *Botrytis cinerea* at 3 days after inoculation. The data are the mean ± SE (*n* >36). Leaves of WT and mutant plants were drop-inoculated with *B. cinerea* spores (2.5 × $10^5$ spores/mL). The experiment was repeated three times, with similar results.

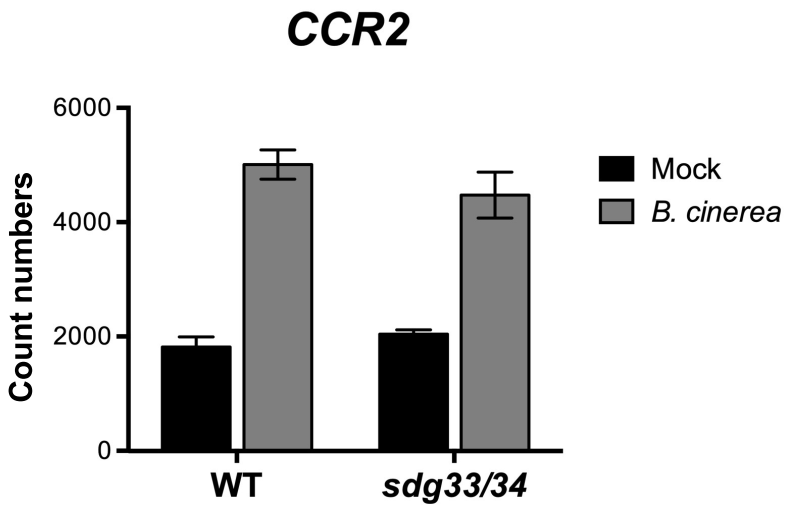


**Fig. S3. Tomato *CAROTENOID ISOMERASE2* expression is independent of SDG33 and SDG34.**
Expression of *CCR2* gene in mock- or *Botrytis cinerea* inoculated WT and *sdg33sdg34* based on RNA-seq transcript count data. Error bars indicate the standard deviation of three libraries.

**Fig. S4. Transcriptome analysis in mock and *Botrytis cinerea* inoculated tomato leaf tissues.**
**(a)** Principal component analysis (PCA) of the normalized RNA-seq data; and **(b)** Gene regulatory network of the 179 genes regulated by genotype. In this network, the triangle-shaped green nodes show transcription factors (TFs). **(c)** A network view of enriched biological processes from the generated network in (**b**).

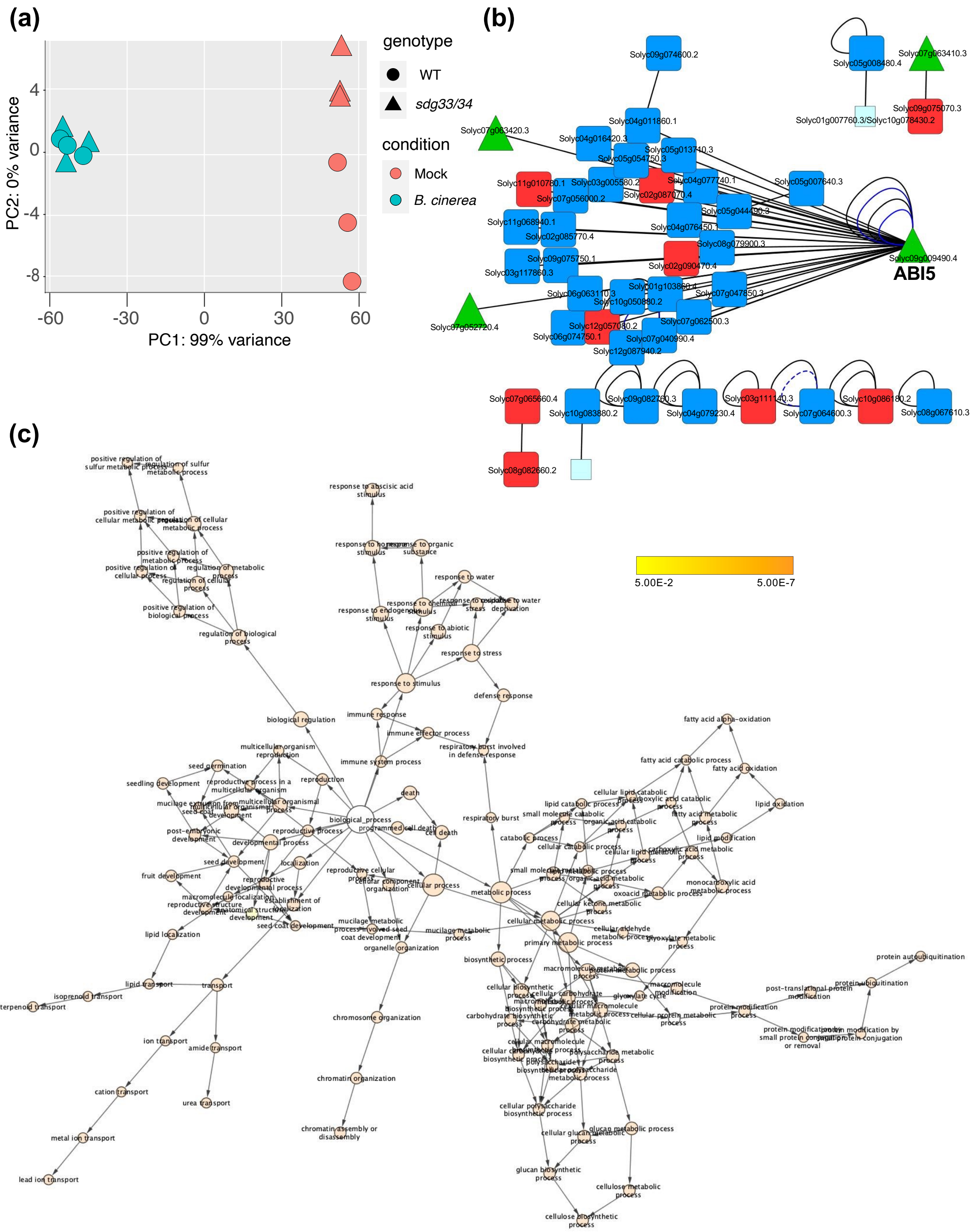


**Fig. S4. Transcriptome analysis in mock and *Botrytis cinerea* inoculated tomato leaf tissues.**

**(a)** Principal component analysis (PCA) of the normalized RNA-seq data; and **(b)** Gene regulatory network of the 179 genes regulated by genotype. In this network, the triangle-shaped green nodes show transcription factors (TFs). **(c)** A network view of enriched biological processes from the generated network in (**b**).

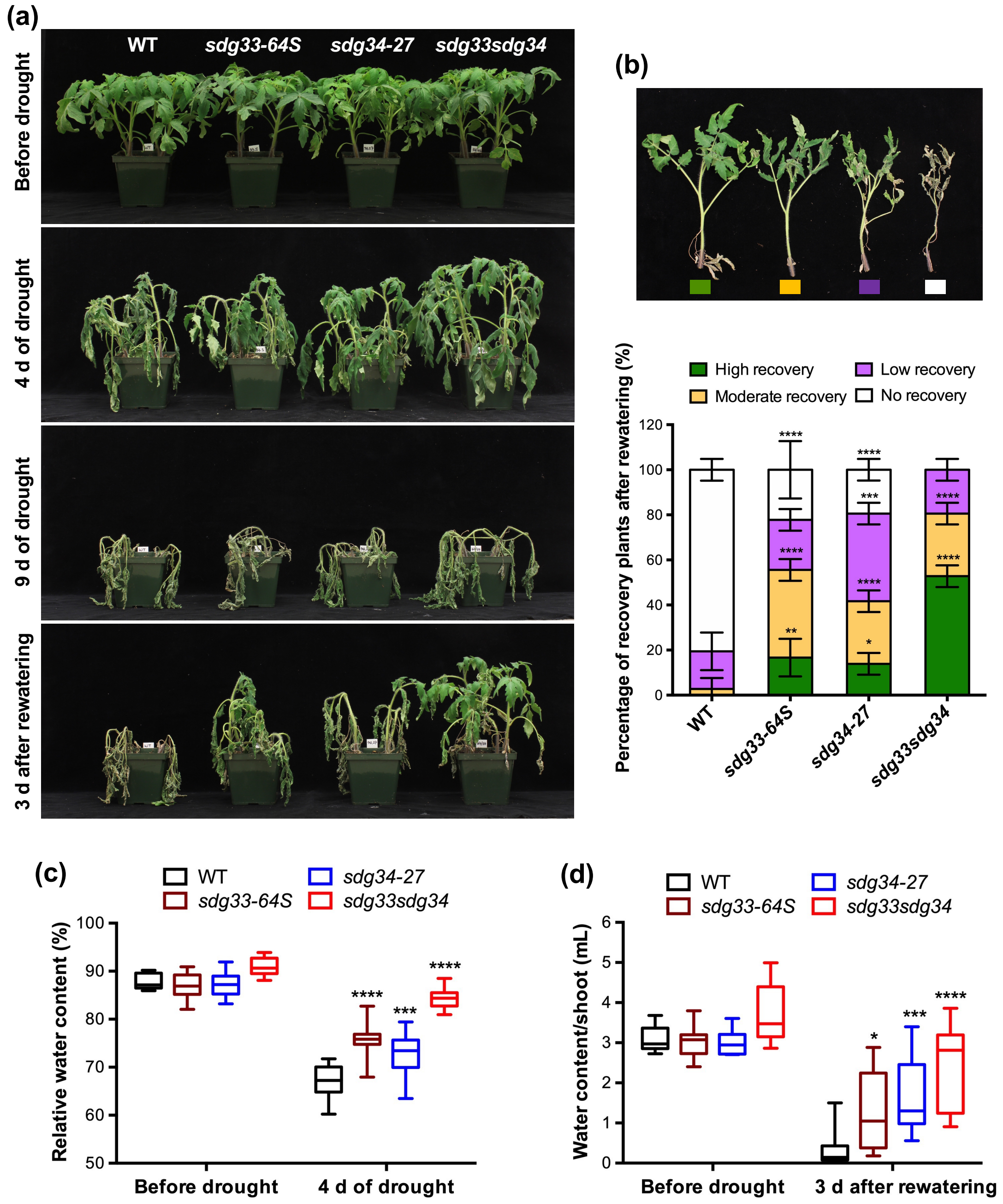


**Fig. S5. The tomato *sdg33* and *sdg34* single and double mutants are tolerant to drought stress in tomato. (A)** Growth and survival of plants before and after drought stress. Four week-old plants were subjected to drought treatment. **(B)** Relative Water Content at day 4 of water stress. The data shown are the mean ± SE ($n = 10$). **(C)** Survival rates and extent of recovery after re-watering of drought stressed plants. The data shown are the mean ± SE ($n = 36$). Data shown is a representation of experiments repeated at least 3 times. Asterisk indicates significant differences ($p < 0.0001$) between the WT and each genotype according to Tukey's multiple comparisons test. **(D)** Shoot water content measured at 3 days after re-watering. The data shown are the mean ± SE ($n = 36$). Asterisks indicate significant differences (*$p < 0.05$, **$p < 0.01$, ***$p < 0.001$, and ****$p < 0.0001$) between the WT and each mutant line counterpart at the same treatment conditions based on Tukey's multiple comparisons test.
